# Viral infection patterns in ants are affected by colony structure and phylogenetic lineage

**DOI:** 10.64898/2026.08.27.747284

**Authors:** Konu Mauno, Chowdhury Riaz Murshed, Abril Sílvia, Cremer Sylvia, Giannetti Daniele, Donato A. Grasso, Helanterä Heikki, Kato Mitsuho, Orivel Jérôme, Ran Hao, Robb Jennifer, Schifani Enrico, Birgit C. Schlick-Steiner, Seppä Perttu, Shimoji Hiroyuki, Florian M. Steiner, Strahodinksy Florian, Trigos-Peral Gema, Tsuji Kazuki, Zijun Xiong, Lequime Sebastian, Viljakainen Lumi

## Abstract

Across ant species, there are differences in how their societies are structured. Single-queened (monogynous) societies only have one reproducing queen in the colony, and new queens disperse and start colonies independently. In multiple-queened (polygynous) societies, the colony instead can contain several reproductive queens, and newborn queens often remain and reproduce within their natal colony. As a result, polygynous societies are comparatively larger, more genetically diverse, and can span large areas through several interconnected nests, whereas monogynous societies are typically smaller in scale. In this study, we investigated how these different social structures, as well as their phylogenetic lineage, affect the diversity (number of virus species per ant sample) and abundance (number of viral sequences per sample) of viruses in ants. We produced pooled RNA sequence libraries from 15 ant species, representing both monogynous and polygynous social structures, and the two largest ant subfamilies: Formicinae and Myrmicinae, with each library containing the RNA of up to 400 individual worker ants from a single population. We identified 168 virus species in total, of which 152 species were new to science. Out of these 168 viruses, 59 were active viruses based on the host immune response. We observed that polygynous ant species harbor a higher diversity of viruses and also tend to have higher virus abundance compared to monogynous species. Also, the ant subfamily Myrmicinae had a higher virus diversity than Formicinae. These findings highlight how social structure and evolutionary history shape viral diversity in ants.

## Introduction

Eusocial insects (ants, termites, and some bees and wasps) are known for their organized social behavior, where tasks are divided between different castes. Only queens produce offspring, while workers, which are typically sterile, collect resources, clean and maintain the colony, and take care of the offspring. The ancestral state of social colony organization in ants is monogyny, where only one queen reproduces in a colony (Boulay et al. 2014, Hughes et al., 2008a, Hughes et al., 2008b). New queens produced in monogynous colonies disperse by flight from their natal colony and found a new colony where they begin to produce offspring. Polygyny, where several queens reproduce in the same colony, has evolved independently in several ant lineages (Boomsma 2009, Hughes et al. 2008, Boulay et al. 2014). In polygynous societies, new queens are often allowed to remain and reproduce in their natal colony, either in the original nest or by founding a satellite nest connected to the other nests of the colony via a path network, and they can share workers (Sundström et al. 2005).

Ants are in constant interaction with pathogens originating from the environments they move in and the food they forage for, and may bring them into their nests. Ant nests themselves are a favorable environment for pathogens to thrive and spread. The nests have stable climate conditions, and the ants live in close proximity to each other, sharing resources and thus risking the spread of pathogens among nestmates (Boomsma et al. 2005, Fürst et al. 2014, French & Holmes 2020). To mitigate the risk of infections in these conditions, the ants exhibit hygiene behaviors, such as cleaning and grooming of fellow workers and bringing waste and dead nestmates out of the nest (Hughes et al. 2002, Rosengaus et al. 1998, Theis et al. 2015, Pull et al. 2018, Rothenbuhler 1964, Tragust et al. 2013, Avanzi et al., 2024, da Silva et al., 2025).

The colony structure of social insects will greatly affect the likelihood of pathogens they encounter, how pathogens spread between nests, and the disease resistance of the colony. Single-queen colonies are typically (excepting leafcutter and driver ants) smaller than multi-queen colonies. The latter therefore often span a larger territory and often have a more open colony structure with higher exchange of individuals between nests. Polygynous colonies are therefore more likely to pick up pathogens from the environment, which could then readily spread between neighboring nests. However, since polygynous colonies are genetically more diverse than the single-queen offspring in monogynous colonies, they are less likely to be eradicated by an incoming pathogen, and thus may be able to tolerate a higher number of infections in their nests (Sherman et al. 1988). We therefore expect that pathogen load and diversity, when measured in surviving field colonies, should be higher in polygynous species, a finding that was recently supported within species in fire ants (Brahma et al., 2022).

Ants are known to harbor many viruses, most of which are specific to one species (Viljakainen et al. 2023, Porter et al. 2013). However, there are also more generalized viruses that infect a wider range of insects, including ants. For example, Black queen cell virus, Deformed wing virus, and Kashmir bee virus were first found in and are usually associated with honey bees, but they are known to infect ants as well, especially if the ant colony is located near a bee colony (Lester et al. 2019, Tiritelli et al. 2024, 2025). In total, almost 3900 ant-associated viruses have been identified. Flynn and Moreau (2024) identified 3710 novel viruses from 35 ant species, while Zueva et al. (2025) reviewed 93 articles and found 187 virus species in 54 ant species, of which 66 are suggested to be replicating (Flynn and Moreau (2024) was published just months before Zueva et al. (2025) and therefore was not cited in the review). The way the pathogens affect the ant phenotype remains poorly understood, and often viruses that are proven to be replicating in ants cause no visible symptoms. Some of the few known symptoms range from decreased queen weight and egg production to impaired mobility and decreased ability to forage and compete against other species, and ultimately death (Lester et al. 2019, Zueva et al. 2025). Pathogens are also thought to be involved in rapid population collapses observed in invasive ant populations (Lester & Gruber, 2016)

There are currently 14,482 known ant species in the world (antcat.org), but viruses and their pathogenicity have been studied mostly on harmful invasive species, like the Argentine ant (*Linepithema humile*) and the Red imported fire ant (*Solenopsis invicta*) (Baty et al. 2020; Viljakainen et al. 2018; Gruber et al. 2017, Manfredini et al. 2016, Valles & Hashimoto 2009). However, since ants are ecologically very important insects and they can be found almost everywhere in the world, their associated viruses may play a more significant role in ecosystems than currently recognized. Understanding ant viruses and their effects can also offer implications that extend beyond ant biology, for example, in chemical-free pest control. Some known viruses with the potential to be used as pest control for ants and termites have been identified, though there are currently no commercially available products (Zueva et al. 2025).

RNA sequencing has been utilized to detect and recognize viruses in different host species, and with this method, thousands of new RNA viruses have been found in invertebrates (Shi et al. 2016, Webster et al. 2015, Viljakainen et al. 2023, Flynn & Moreau 2024). To identify active infections, techniques that sequence both long and short RNAs have been combined, so that small interfering RNAs (siRNAs) induced by host immune responses can be included in the analyses (Webster et al. 2015, Viljakainen et al. 2023). The process involving siRNAs, called RNA interference, both regulates gene expression and prevents virus infections using short RNA fragments. Viral double-stranded RNA is broken down, and the pieces are used as a template to destroy viral RNAs (Meister & Tuschl, 2004). By connecting these viral RNA fragments to the found virus genomes, the host immune response, and its intensity against the viruses can be detected.

This study concentrates on characterizing the diversity of RNA viruses as the number of virus species per ant sample, and the virus abundance as the number of viral sequences per sample, in fifteen ant species by using RNA sequencing data. We expect that the ant species’ virome will be affected both by their natural history, reflected by their phylogenetic lineage and their social colony structure, having either one or multiple queens and thus representing either small, closer colonies of highly related individuals, or larger, more open colonies with a higher genetic diversity. We expect in particular that polygynous colonies may harbour a greater viral diversity and abundance since their social structure both favours transmission between nests and higher survival chances upon infection.

## Materials and methods

### Ant sample collection

This study was conducted with 15 ant species across the world (Table 1). The species were selected as closely related pairs, one monogynous and the other polygynous; however, for two of the species (*Crematogaster levior* and *Temnothorax nylanderi*), a corresponding counterpart could not be obtained. Additionally, two of the species (*Myrmica ruginodis* and *My. scabrinodis*) are known to have both monogynous and polygynous populations, so they were divided into a separate “socially polymorphic” category. *Formica pratensis* is also known to have polygynous populations, though the populations in Finland, where they were collected, are predominantly monogynous, thus it was categorised as such. The studied ants were collected from various locations worldwide. For each species, samples were collected from a single population, comprising 5-20 worker ants each from 15-20 nests, yielding 290-400 worker individuals per species (Table 1). The ants were collected directly in RNA*later* (ThermoFisher) and stored at - 20 °C before RNA extraction.

**Table 1.** Origin of the ant samples and numbers used in the RNA extraction.

|  | Country | Number of nests sampled | Number of ants extracted | Ants per extraction batch |
| --- | --- | --- | --- | --- |
| *Crematogaster levior* | French Guiana | 20 | 400 | 10-20 |
| *Formica aquilonia* | Finland | 20 | 400 | 5 |
| *Formica pratensis* | Finland | 20 | 400 | 5 |
| *Lasius neglectus* | Germany | 20 | 400 | 10-20 |
| *Lasius niger* | Austria | 20 | 400 | 10-20 |
| *Messor barbarus* | Spain | 20 | 400 | 10 |
| *Messor structor* | Austria | 20 | 368 | 5-10 |
| *Myrmica rubra* | Poland | 20 | 400 | 10 |
| *Myrmica ruginodis* | Finland | 20 | 385 | 11-20 |
| *Myrmica scabrinodis* | Poland | 20 | 400 | 10 |
| *Pheidole fervida* | Japan | 20 | 400 | 20 |
| *Pheidole megacephala* | Japan | 20 | 400 | 20 |
| *Temnothorax nylanderi* | Italy | 15 | 290 | 10-20 |
| *Tetramorium bicarinatum* | China | 20 | 400 | 10-20 |
| *Tetramorium caespitum* | Finland | 20 | 400 | 20 |

### RNA extraction

The RNA extraction was performed using the mirVana™ miRNA Isolation Kit (ThermoFisher) and following the Total RNA Isolation Procedure protocol, which simultaneously isolates both long and short RNA molecules. For each species, the RNA was extracted in batches, the size of which depended on the size of the individual ants (Table 1), and the RNA quality and quantity in each batch was determined using the RNA 6000 Pico Kit on a 2100 Bioanalyzer system. Separate extractions were then pooled into a single large sample for each species, with equal quantities of RNA from each extraction. From half of this sample, the long RNAs were sequenced (fragment size >350 nt, directional library preparation, rRNA removal, Illumina NovaSeq paired-end 150 bp long reads), and from the other half, the short RNAs (sRNAs, fragment size <50 bp, small RNA library preparation, Illumina NovaSeq single-end 50 bp long reads) were sequenced at Novogene.

### Virus genome assembly

The long, 150 bp RNA reads were used to determine the virus species present in the ant sample and their relative abundance, through assembly into complete virus genomes. The long RNA data were first run through Lazypipe (version 2.1) (Plyusnin et. al. 2020) and then CAP3 (Huang & Madan, 1999) to assemble the RNA reads into longer contigs. Lazypipe utilizes the host genome for each ant species in addition to the RNA sequence data. The program compares RNA reads against the host genome and excludes all matches, leaving only foreign RNA from viruses, bacteria, and other eukaryotes. The host species’ genome was used whenever it was available, and when unavailable, the reference genome of the closest relative was used (Table 2). All but one of the genomes (the hybrid *Formica aquilonia × F. polyctena* genome, GenBank: CAJQTV000000000.1) were provided as early access by the Global Ant Genomics Alliance (GAGA) (Vizueta et al. 2025). Lazypipe categorizes contigs into three groups using SANSparallel and BLAST to search for similar sequences from NCBI nt, RefSeq, and GenBank databases: eukaryotes, bacteria, and viruses, and the contigs classified as virus-derived were selected for further analysis. Due to the high mutation rate of viruses, many variants of the same virus were found in the sample. They were combined into one representative sequence using CAP3, which allows more sequence variation than Lazypipe and, therefore, can better combine similar sequences that Lazypipe would otherwise separate into two species. These representative sequences of the virus genomes were then screened with a custom script to remove sequences shorter than 1000 nt. Genome sizes in RNA viruses can range from 2-41 kb (Lauber et al. 2013, Ferron et al. 2021), so by removing short contigs, the analyses could be targeted to complete or close-to-complete virus genomes.

**Table 2.**
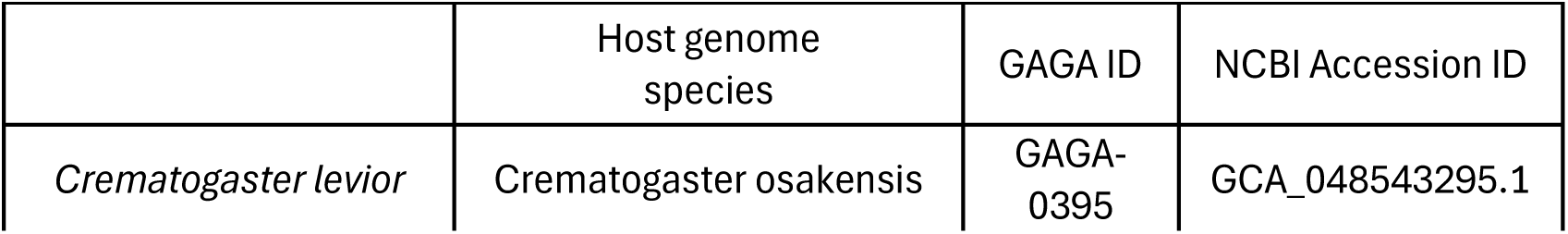
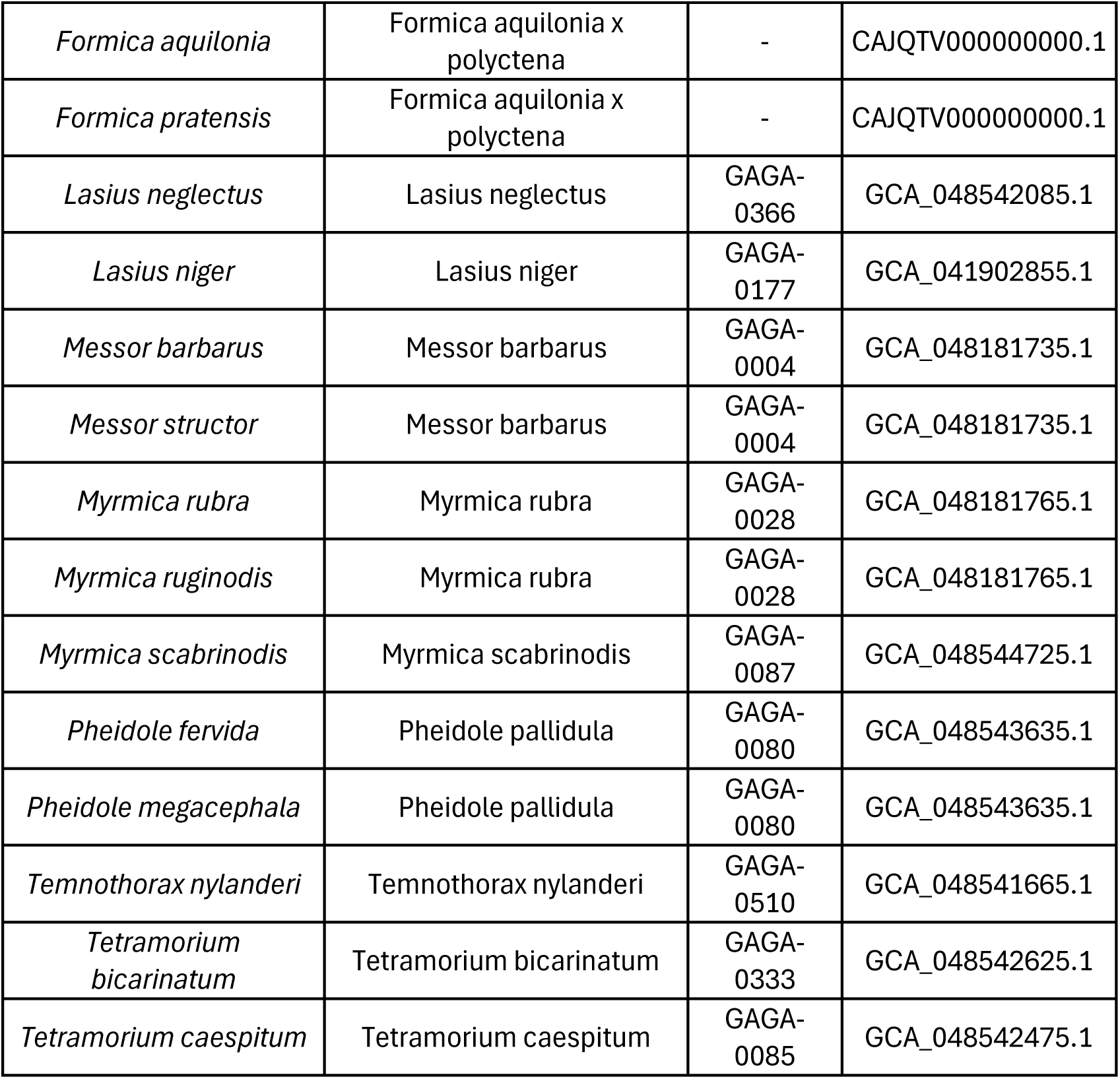
Host genome used in Lazypipe.

### Virus abundance estimation

To estimate the abundance of viruses in the ant samples and compare the results across different species, the RPKM value (Reads Per Kilobase per Million mapped reads) was calculated. Three numbers are needed for the calculation: the number of mapped reads (how many reads mapped to this virus genome), a scaling factor (the total number of quality-filtered RNA reads in the dataset), and the virus genome length (in kilobases). The scaling factor was obtained by quality-filtering the RNA raw reads with Trimmomatic (Bolger et al., 2014). To obtain the number of mapped reads and contig length, the RNA sequence reads were mapped against the virus genomes with a custom script that utilizes BWA and samtools (Li, 2013 & Li et al., 2009).

### Assessing RNAi-based immune response against viruses

sRNAs were used to detect the RNA interference (RNAi) response that the ants induce against viruses. RNAi is based on double-stranded RNA that is cleaved into 21-22 nt segments (Zotti et al., 2026) in the cytoplasm by the Dicer protein and loaded into the Argonaute protein, forming RISC (RNA-induced silencing complex), which uses the segments as a template to detect similar sequences. The detected RNAs are then cleaved and degraded, halting viral replication (Wilson and Doudna, 2013). Based on the relative amount of the degraded short 21-22 nt RNA segments, we can infer an active immune response against RNA viruses. To reveal the immune response, the RPKM values for the sRNAs were calculated and their size distribution assessed. First, the adapter sequences used for sequencing were removed using the sRNA_clean script in VirusDetect (Zheng et al., 2017). The cleaned sRNA data were then mapped against the virus genomes. The mapping script utilizes Bowtie (Langmead, 2010) and samtools, providing a size distribution of sRNA sequences. The results include the genome (contig) length, the number of mapped reads, and the number of cleaned reads (scaling factor), which were used for the sRNA RPKM calculation. Then, the R package viRome (Watson et al., 2013) was used to visualize the size distribution of sRNA data and reveal RNAi response against viruses. The peak of 21-22 nt-long RNA segments, in addition to an RPKM value >10, was used as an indicator of an immune response to the virus and therefore as a sign of an active virus infection.

### Annotation of the virus genomes

To focus further analyses on the most abundant viruses, only the virus genomes with the long and/or short RNA RPKM value above 10 were chosen for annotation. The annotation was conducted with NCBI ORF finder (https://www.ncbi.nlm.nih.gov/orffinder/), with a minimum ORF length set to 300 nt. Each ORF was run through a BLAST search (Reference proteins (refseq_protein) as the BLAST database) to find out which gene the ORF represents. If the ORF contained an RNA-dependent RNA polymerase (RdRp) protein-encoding gene, the NCBI Conserved Domain Database (CDD) (Wang et al., 2023) was searched, and the exact location of the protein in the amino acid sequence was marked down for phylogenetic classification based on RdRp. In some cases, there was no CDD available for the RdRp, so the BLAST search’s alignment graphics and the GenPept database were utilized to find the RdRp location.

### Phylogenetic analysis

To taxonomically classify the viruses and determine their phylogenetic relationships, the annotated virus contigs were compared with known viruses, called here the control group. A family-level phylogenetic tree of the control group viruses was constructed following Wolf et al. (2018). The ICTV’s Virus Metadata Resources Excel sheet (version VMR_MSL39_v2.xlsx, downloaded 2.10.2024) (https://ictv.global/vmr) was downloaded, and a representative species from each RNA virus family was picked, totaling 138 virus species, thus representing 138 viral families. Species that lacked family or higher taxonomic classification were also included, as long as they were classified into a known phylum. The amino acid sequences of the RdRp genes from each of these control group viruses were then extracted.

Constructing a single tree that included all viruses and the control group was not possible due to substantial sequence differences. The data had to be divided into smaller chunks for easier handling. In their article, Wolf et. al. (2018) concluded that RNA viruses can be divided into five distinct clades. The control group was divided into these five subgroups similarly. The trees in Wolf et. al. (2018) were used to select the corresponding viruses from the control group data. Once all the viruses mentioned in the article were selected, it was concluded that these subgroups exactly match the phylum of the viruses. With this information, the control group data could be split into seven subgroups representing all seven viral phyla (two new phyla have been characterized after 2018): Lenarviricota (14 families), Pisuviricota (36 families), Kitrinoviricota (21 families), Duplornaviricota (22 families), Negarnaviricota (37 families), Ambiviricota (4 families), and Artverviricota (4 families). In addition, an eighth subgroup was created that contained every known genus of the order Picornavirales (105 genera), which falls into the phylum Pisuviricota. This was done because it was predicted that a large proportion of viruses would fall into this order (Viljakainen et al. 2023), making it easier to determine the virus species later.

The final trees were constructed in several iterations. The 198 virus RdRp amino acid sequences were aligned against each control subgroup. First, the sequences were aligned with MAFFT (Katoh & Standley, 2013). with the E-INS-I algorithm. Next, low-quality amino acid residues were masked at the 10% fractile using Guidance 2 (Sela et. al. 2015), and finally, phylogenetic trees were reconstructed with IQTree (version iqtree-2.3.6-Linux-intel) (Nguyen et al. 2014) (parameters: -m MFP -mtree -nt AUTO -B 1000 -bnni) and visualized with iTOL (Letunic & Bork, 2024). Once the first trees were reconstructed, it was observed which viruses fell within each control subgroup. Those that were uncertain were extracted from the original batch and re-run against each control subgroup to obtain more precise results with smaller batches.

Some viruses were assigned to two or more control subgroups. These overlapping viruses were resolved by observing the involved subgroup trees to see whether the overlapping viruses formed a monophyletic group within the tree, whether the control viruses were included within this monophyletic group or outside of it, and how strong the bootstrap values were within this monophyletic group compared to the branch connecting it to the rest of the tree. Once all the viruses were classified into control subgroups, the final phylogenetic trees for each control subgroup were reconstructed. The final phylogenetic trees contained only the control subgroup and the viruses that were classified into this subgroup, yielding the most accurate trees.

### Virus species classification

The ICTV Report Chapters by Genome (ictv.global/report/genome) for species demarcation criteria were used for virus species classification. If this source did not have the necessary information, the ICTV Approved Proposals (ictv.global/files/proposals/approved) were consulted for additional information (E. J. Lefkowitz, personal communication, 17.6.2024). For each virus species, the closest control subgroup virus and its taxonomic family were searched from the phylogenetic trees. This family was then searched from the Report Chapters, and according to the genus and species demarcation criteria, the virus was classified as either a new or known species. Most commonly, the demarcation criteria demanded a certain degree of similarity for the RdRp gene; for example, less than 90% identity to the closest match in the database would be considered a new species, but some virus families had different criteria, like the identity of the capsid protein or the entire genome sequence. The viruses were named according to the ant species and the closest related virus family name, followed by a number representing the order of discovery. For example, “Myrmica rubra phenui-like virus 1” was a virus found in the ant *My. rubra*, and the virus is related to the Phenuiviridae family and was the first of this kind found. In cases where the closest related family did not have demarcation criteria available or the closest related group was uncertain, several closely related families were consulted to form consensus criteria that would most likely apply to this virus. In cases where it was unclear whether a virus should be classified as a new species or a known species, it was named a new virus because, presumably, it would be easier to later combine two species into one than to divide one species into two. The names were finalized and confirmed after finding possible duplicates using BLAST search.

### Reciprocal BLAST search

A reciprocal BLAST search, where the ant virus genomes were compared against each other, was done to find out if there were any viruses similar enough that they could be considered the same species. The BLAST search was performed for each subgroup separately. The matching sequences were then reviewed to determine whether the alignment length and sequence identity were sufficiently similar. When the sequences were of similar length and the whole alignment identity was at least 95%, they were declared the same species, and the names were updated to match. When the identity was around 80%, a manual check was performed to determine whether the demarcation criteria for these viruses justified treating them as the same species. For example, in these cases, the RdRp aa-sequence identity was often much higher than the whole-genome identity, leading them to be considered the same species. In cases where the identity was 70% or below, the alignment length also became very short, spanning less than half the length of the virus genomes. These cases were judged to be separate species. When the same virus was found in two or more ant species, the virus name was updated to match the lowest common taxonomic level of the ant species. For example, Myrmicinae picorna-like virus 1 was found in four different ant species: *Messor barbarus, My. rubra, My. ruginodis,* and *My. scabrinodis*. These four ant species belong to the Myrmicinae subfamily.

### Identification of viruses derived from the sRNA data

VirusDetect (Zhen et. al. 2017) is a program used to identify viruses from sRNA data. It assembles sRNA sequences into longer contigs and compares them against a database (GenBank gbvrl) of known viruses, much like Lazypipe does with long RNA sequences. It was used to visualize how the host ant’s immune response degrades double-stranded viral RNA into smaller fragments (range 18-33 nt) and to identify viruses that the host’s immune response might have already destroyed, leaving only traces of sRNA rather than complete virus genomes.

VirusDetect was run separately for each of the 15 ant species’ sRNA data, and a total of 60 known virus sequences were found that matched the sRNA contigs. However, many of these alignments contained either very short known virus sequences, or there were only a few sRNA contigs aligned with the known virus sequence, or the sRNA contigs covered only a very limited range of the known virus sequence. Therefore, low-quality alignments were excluded using the following criteria: The known virus sequence had to be longer than 1000 nt, sRNA contigs had to cover more than 10% of the known virus sequence, and there had to be more than 5 sRNA contigs matching the known virus sequence. Screening the alignments using these criteria resulted in 21 high-quality alignments that were selected for further analysis.

The 21 virus sequences obtained from the high-quality alignments were used as BLAST queries against all virus genomes identified from the long RNA data. When a query sequence matched a long RNA-derived virus genome from the same ant species as the sRNA contigs, the corresponding sRNA contigs were assigned to that virus genome. This indicated that the sRNA contigs originated from the matching long RNA-derived virus genome. If a search sequence did not match any long RNA-derived virus genome, it indicated that this virus could only be found in the sRNA data and was a virus that the host ant’s immune system had already destroyed to the point where it could no longer be detected in the long RNA data. Alternatively, the sRNAs could have originated from endogenous viruses that Lazypipe screened out because they match the host genome.

## Statistical analysis

All statistical tests were conducted in R (v4.5.2) (R Core Team, 2021), and statistical significance was evaluated at an alpha level of 0.05 for all analyses. One goal of this study was to determine if the virus diversity and total viral abundance differ between monogynous and polygynous ant species. Due to sample sizes, the statistical test was conducted with only two groups, monogynous (n=6) and polygynous (n=7), using the Wilcoxon rank-sum test, leaving out the socially polymorphic species (n=2). To compare the virus diversity between monogynous and polygynous ants, the number of unique virus species in each ant species was calculated (Table 3). To compare the total viral abundances in monogynous and polygynous ants, the RPKM values for each virus species found in the ant were combined to get the value for total viral abundance in each ant species.

**Table 3.** Number of unique viruses in each ant species.

| Monogynous species | Unique viruses | Polymorphic species | Unique viruses | Polygynous species | Unique viruses |
| --- | --- | --- | --- | --- | --- |
| *Formica pratensis* | 5 | *Myrmica scabrinodis* | 14 | *Formica aquilonia* | 5 |
| *Lasius niger* | 5 | *Myrmica ruginodis* | 18 | *Lasius neglectus* | 3 |
| *Messor barbarus* | 6 |  |  | *Messor structor* | 13 |
| *Tetramorium caespitum* | 7 |  |  | *Myrmica rubra* | 25 |
| *Pheidole fervida* | 15 |  |  | *Tetramorium bicarinatum* | 13 |
| *Temnothorax nylanderii* | 4 |  |  | *Pheidole megacephala* | 37 |
|  |  |  |  | *Crematogaster levior* | 16 |
| Total number | 42 | Total number | 32 | Total number | 112 |
| Per species average | **7** | Per species average | **16** | Per species average | **16** |

Additionally, a second set of statistical tests was conducted where the aspect of closely related species pairs of the ants was taken into account. Here, the tests were conducted with a total of ten ant species divided into five pairs: *F. pratensis* and *F. aquilonia, Lasius niger* and *L. neglectus*, *Me. barbarus* and *Me. structor*, *Pheidole fervida* and *P. megacephala*, and *Tetramorium caespitum* and *Tet. bicarinatum* (note that the *Tetramorium* species pair is not particularly closely related). The remaining five species, *C. levior, My. scabrinodis, My. ruginodis, My. rubra*, and *Tem. nylanderi* were excluded from these tests.

As with the first set of tests, the species pairs were first analysed on the account of virus species diversity using the negative binomial generalized linear mixed model (GLMM) (glmmTMB, v1.1.14, Brooks et al. 2017), with social structure (monogynous vs. polygynous) as the fixed effect and species pairs as the random intercept. Total viral abundance per ant species was analysed using the linear mixed-effect model on log-transformed data (lme4, v2.0-1, Bates et al. 2015). Social structure was used as the fixed effect, and species pairs as the random intercept. Viral abundance at the level of individual viruses (RPKM value for each individual virus) was analysed using the linear mixed-effect model with species as a random effect (lme4, v2.0-1, Bates et al. 2015). A paired Wilcoxon signed-rank test was used as a robustness test. The relationship between the virus species diversity and total viral abundance was determined using Spearman’s rank correlation test.

We noticed that the ants in the subfamily Formicinae appeared to have much fewer virus species compared to the ants in the subfamily Myrmicinae regardless of the social structure of the species. Therefore, we tested whether the subfamily has an effect on virus diversity. Differences in viral diversity between the ant subfamilies Formicinae (n = 4) and Myrmicinae (n = 11) were assessed using the Wilcoxon rank-sum test. Viral diversity was defined as the total number of virus species detected in each ant species.

To determine whether viral community composition differed between ant subfamilies, a permutational multivariate analysis of variance (PERMANOVA) was performed using Bray–Curtis dissimilarities in vegan v2.7-5 (Oksanen et al., 2026) (adonis2, 9,999 permutations). The analysis was based on the number of virus species assigned to each viral phylum (Lenarviricota, Pisuviricota, Kitrinoviricota, Duplornaviricota and Negarnaviricota). Picornavirales was included within Pisuviricota for this analysis because it represents a lineage within that phylum. Because visual inspection suggested that Pisuviricota viruses dominated the observed differences between subfamilies, an additional exact Wilcoxon rank-sum test was performed to compare the number of Pisuviricota virus species between Formicinae and Myrmicinae.

Bray–Curtis dissimilarities were visualized using non-metric multidimensional scaling (NMDS). To verify that significant PERMANOVA results were not caused by differences in within-group dispersion, homogeneity of multivariate dispersions was tested using *betadisper* followed by a permutation test.

## Data visualization

Graphs depicting virus diversity, genome structure, and statistical results were created in R (v4.5.2) using the packages ggplot2 (v4.0.2, Wickham, 2011), ggrepel (v0.9.7, Slowikowski, 2026), circlize (v0.4.18, Gu et al. 2014), dplyr (v1.2.0, Wickham et al. 2026), readr (v2.2.0, Wickham et al. 2026), and tidyverse (v2.0.0, Wickham et al., 2019). Figure depicting the sRNA identity compared to complete virus genomes was assembled using the imagery created by VirusDetect (Zhen et. al. 2017).

## Results

### Virus genome assembly

A total of 5025 virus genomes were initially assembled, ranging from 29 genomes derived from *L. neglectus* to 1039 genomes derived from *P. megacephala* (Table 4). After merging highly similar virus variants using the sequence assembly program CAP3, the number of virus genomes dropped from 5025 to 3470, ranging from 24 genomes for *L. neglectus* to 502 genomes for *P. megacephala* (Table 4). Finally, the short, less than 1000 nt long sequences were removed. This reduced the total number of virus genomes to 1297 (26% of the initially discovered 5025 viruses), ranging from 13 genomes for *L. neglectus* to 192 genomes for *P. megacephala* (Table 4).

**Table 4.** The number of RNA sequences after each step of analysis. The final number of viruses is 168 (154 novel viruses and 14 known viruses) after the duplicate viruses were combined from the initial 198 viruses. The total number of unique virus species (186) differ from the final number of viruses (168) since some viruses were found from two or more ant species, and therefore those viruses appear in the count several times.

|  | Raw data read pairs | Read pairs after trimmomatic filtering | Lazypipe | Cap 3 | Sequences longer than 1000 nt | RPKM & siRNA RPKM > 10 | RdRp | Novel viruses | Known Viruses | Final Novel Viruses | Final Known Viruses | 1. Lenarviricota | 2. Pisuviricota | 2.1. Picornavirales | 3. Kitrinoviricota | 4. Duplornaviricota | 5. Negarnaviricota | Total number of unique virus species in each ant species |
| --- | --- | --- | --- | --- | --- | --- | --- | --- | --- | --- | --- | --- | --- | --- | --- | --- | --- | --- |
| *Crematogaster levior* | 37431111 | 36525998 | 754 | 476 | 174 | 36 | 22 | 21 | 1 |  |  | 0 | 3 | 10 | 1 | 1 | 1 | 16 |
| *Formica aquilonia* | 40794729 | 39889024 | 57 | 29 | 17 | 8 | 5 | 5 | 0 |  |  | 0 | 1 | 1 | 0 | 1 | 2 | 5 |
| *Formica pratensis* | 37677689 | 36913018 | 214 | 128 | 69 | 25 | 5 | 5 | 0 |  |  | 0 | 2 | 1 | 0 | 1 | 1 | 5 |
| *Lasius neglectus* | 35341673 | 34515263 | 29 | 24 | 13 | 5 | 3 | 1 | 2 |  |  | 0 | 0 | 1 | 0 | 1 | 1 | 3 |
| *Lasius niger* | 45611935 | 44689068 | 79 | 33 | 14 | 5 | 5 | 3 | 2 |  |  | 0 | 0 | 5 | 0 | 0 | 0 | 5 |
| *Messor barbarus* | 37729964 | 36957554 | 363 | 307 | 98 | 9 | 6 | 4 | 2 |  |  | 0 | 2 | 1 | 3 | 0 | 0 | 6 |
| *Messor structor* | 24808884 | 24425625 | 238 | 210 | 74 | 22 | 13 | 12 | 1 |  |  | 1 | 2 | 6 | 2 | 1 | 1 | 13 |
| *Myrmica rubra* | 35685467 | 34823907 | 458 | 319 | 132 | 31 | 26 | 24 | 2 |  |  | 2 | 3 | 13 | 3 | 1 | 3 | 25 |
| *Myrmica ruginodis* | 42502103 | 41719529 | 381 | 271 | 90 | 24 | 19 | 19 | 0 |  |  | 1 | 9 | 7 | 1 | 0 | 0 | 18 |
| *Myrmica scabrinodis* | 31270434 | 30605338 | 236 | 189 | 83 | 28 | 17 | 16 | 1 |  |  | 0 | 2 | 6 | 2 | 2 | 2 | 14 |
| *Pheidole fervida* | 20745005 | 20328225 | 445 | 420 | 142 | 17 | 15 | 14 | 1 |  |  | 0 | 1 | 12 | 2 | 0 | 0 | 15 |
| *Pheidole megacephala* | 33413465 | 32788458 | 1039 | 502 | 192 | 53 | 38 | 34 | 4 |  |  | 2 | 1 | 30 | 4 | 0 | 0 | 37 |
| *Temnothorax nylanderii* | 29718884 | 29208417 | 33 | 33 | 14 | 6 | 4 | 4 | 0 |  |  | 0 | 1 | 2 | 0 | 1 | 0 | 4 |
| *Tetramorium bicarinatum* | 31310864 | 30626499 | 363 | 258 | 97 | 17 | 13 | 13 | 0 |  |  | 0 | 0 | 12 | 1 | 0 | 0 | 13 |
| *Tetramorium caespitum* | 41087647 | 40342435 | 336 | 271 | 88 | 9 | 7 | 7 | 0 |  |  | 0 | 1 | 3 | 0 | 1 | 2 | 7 |
| Total |  |  | 5025 | 3470 | 1297 | 295 | 198 | 182 | 16 | 154 | 14 |  |  |  |  |  |  | 186 |

### Viruses selected for annotation

To focus our analysis of the ant viruses on the most abundant ones, we selected the virus genomes with either long or short RNA RPKM values above 10 for annotation and further analysis (Figure 1). This selection reduced the number of analysed virus contigs from 1297 to 295 (23%, or 6% of the initially discovered 5025 viruses). The number of the remaining virus genomes ranged from 5 for *L. neglectus* to 53 for *P. megacephala* (Table 4).

**Figure 1.**
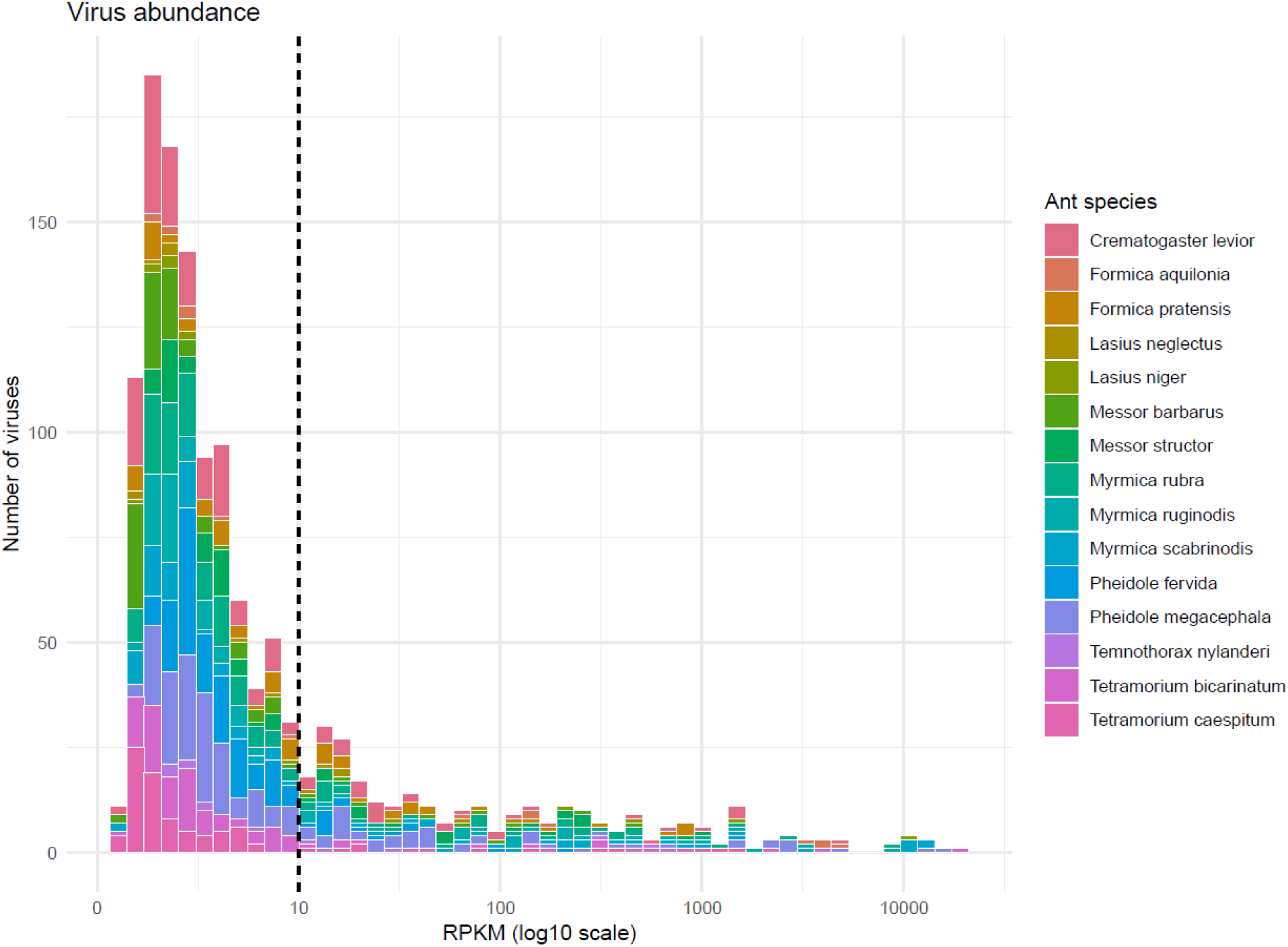
The virus genome RPKM value distribution for all genomes that were longer than 1000 bp (1297 genomes in total) among the 15 ant species. Each virus genome had a separate RPKM value for long RNA and short RNA, but in this graph, only the higher of the two values was selected. There were 295 (23%) virus genomes with either or both RPKM values above 10 and therefore were selected for annotation.

The prediction of ORFs and their annotation revealed that 198 (67%) out of the identified 295 virus genomes contained the RdRp-encoding gene, which was used for the phylogenetic analysis (Supplementary Table 1). The number of selected RdRp-containing virus contigs per species ranged from 3 contigs for *L. neglectus* to 38 contigs for *P. megacephala* (Table 4).

### Phylogenetic analysis

The list of control group viruses used in the phylogenetic analyses contained 138 RNA virus species, each representing a viral family. The control group viruses were divided into seven control subgroups, representing the seven viral phyla, and the number of species per subgroup ranged from 4 to 37, plus the eighth subgroup for the order Picornavirales (105 genera) (see methods). The 198 ant viruses were aligned against each control subgroup to determine which control subgroup the viruses were related to.

In some cases, the virus genomes aligned into two or more control subgroups. For example, all the 19 viruses that aligned well with the Kitrinoviricota control subgroup also aligned well with the Pisuviricota control subgroup. In both cases, the viruses formed a monophyletic group. For the Kitrinoviricota tree, the monophyletic group contained all 19 viruses and all 21 Kitrinoviricota controls. For the Pisuviricota tree, the monophyletic group contained 18 ant viruses (the one remaining aligned somewhere else) and not a single Pisuviricota control. This indicated that the 19 viruses belong to the phylum Kitrinoviricota, and they were excluded from the final Pisuviricota tree. A similar situation occurred with the Duplornaviricota and Negarnaviricota, both of which also aligned with Pisuviricota, but were finally excluded from the Pisuviricota tree.

Bootstrap values were used as directional information when assigning the phylum for viruses. For example, in the Duplornaviricota tree, there were 12 viruses and 22 controls that formed a monophyletic group. However, there was a virus (Myrmica rubra yadokari-like virus 1) that had a bootstrap value of only 8 to its neighboring control. The same virus also aligned into the Pisuviricota tree, where it had a bootstrap value of 100 to its neighboring control. Therefore, the virus was excluded from the final Duplornaviricota tree. Another virus (Myrmica rubra alphatetra-like virus 1) in the Duplornaviricota tree also aligned into the Pisuviricota tree and Kitrinoviricota tree. In the Duplornaviricota tree, the virus had a bootstrap value of 73 to its neighboring virus, in the Pisuviricota tree, the virus had a bootstrap value of 67 to its neighboring virus, and in the Kitrinoviricota tree, the virus had a bootstrap value of 98 to its neighboring control. Therefore, this virus was assigned to Kitrinoviricota, and the Duplornaviricota tree was left with 10 viruses and 22 controls.

Once the virus contigs were confirmed in their phyla, the final trees were reconstructed with only the control groups and the viruses that aligned with them to get the most accurate phylogenies. Lenarviricota tree contained 6 viruses and 14 controls, Pisuviricota tree contained 150 viruses and 36 controls, Kitrinoviricota tree contained 19 viruses and 21 controls, Duplornaviricota tree contained 10 viruses and 22 controls, Negarnaviricota tree contained 13 viruses and 37 controls. The Picornavirales tree contained 121 viruses (note, these viruses were also included in the Pisuviricota tree) and 105 controls.

### Virus species classification

The identified and annotated ant virus species could be classified into five viral phyla: Lenarviricota, Pisuviricota, Kitrinoviricota, Duplornaviricota, and Negarnaviricota. No viruses from the viral phyla Ambiviricota and Artverviricota were found. The classification was carried out starting from the bottom of the tree, going up, since the uppermost viruses were often those that were the furthest away from known controls and often fell outside known families. For each virus, the branches were followed towards the root to find the closest control subgroup virus. The control virus’s taxonomical family was then searched from the ICTV Report Chapters or Approved Proposals to find the demarcation criteria. The virus was then compared, according to the demarcation criteria, to the closest matching known virus found during the annotation process to confirm if the virus is the same species as the annotation match or whether the virus could be classified as a new species and named accordingly.

Below are descriptions of an example virus for each phylum, and other notable cases. Figure 2 shows the visualization of the genome structure for each example virus.

**Figure 2.**
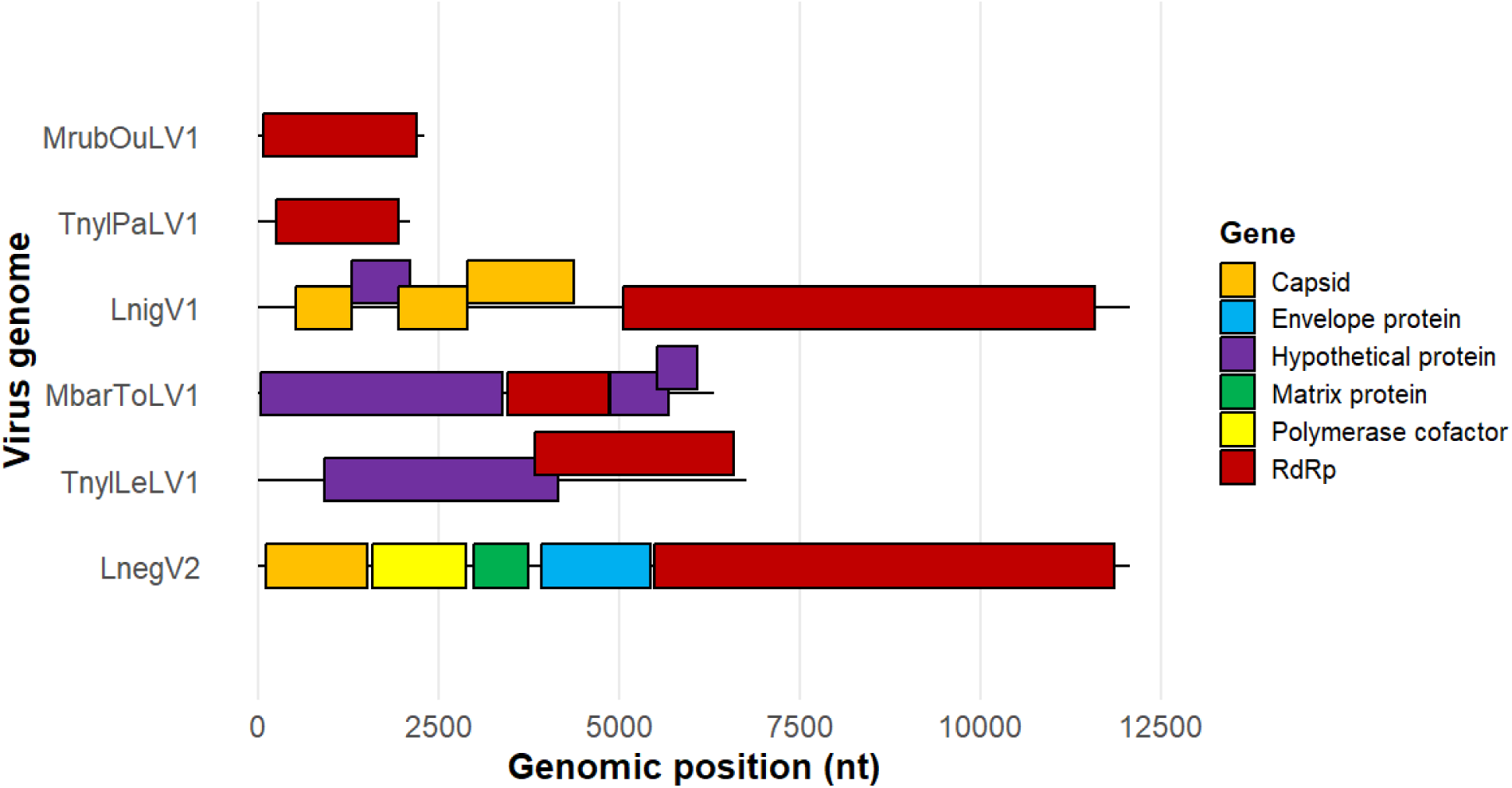
Size distribution and gene locations for each example virus genome. MrubOuLV1 = Myrmica rubra ourmia-like virus 1 (Lenarviricota), TnylPaLV1 = Temnothorax nylanderi partiti-like virus 1 (first half of segmented genome) (Pisuviricota), LnigV1 = Lasius niger virus 1 (Picornavirales), MbarToLV1 = Messor barbarus tobamo-like virus 1 (Kitrinoviricota), TnylLeLV1 = Temnothorax nylanderi leboti-like virus 1 (Duplornaviricota), LnegV2 = Lasius neglectus virus 2 (Negarnaviricota).

### Lenarviricota

Lenarviricota is a phylum of positive single-stranded RNA viruses (Sadiq et al., 2022). Many of the known virus families of this phylum did not have a Report Chapter and therefore had no demarcation criteria. Hence, a prediction had to be made about what they might be according to those families that had the demarcation criteria available. Six ant viruses fell into this phylum (Supplementary Figure 1), of which one is represented here as an example:

Myrmica rubra ourmia-like virus 1. In the final phylogenetic tree, this virus appeared as an outgroup, as if the other viruses and controls were more related to one another than to this virus. However, in the earlier iteration of the phylogenetic tree, this virus paired up with the Botourmiaviridae control, and the closest matching virus found in annotation, Hubei mosquito virus 3, belongs to this family. Therefore, we decided to use the Botourmiaviridae family’s demarcation criteria. Viruses in the Botourmiaviridae family typically infect plants and fungi (Ayllón et al., 2020). They are positive-sense RNA viruses with a mostly 2-5 kb monopartite genome or tripartite genome in the case of the genus *Ourmiavirus* (Ayllón et al., 2020). Myrmica rubra ourmia-like virus 1 had a 2,3 kb long monopartite genome where the ORF containing the RdRp gene is located at 77-2197 nt (Figure 2). This genome size and RdRp location are similar to those of the twelve known genera. The annotated genome was negative-sense, implying that the sequence was a translated messenger RNA instead of an actual genome. According to the genus-level demarcation criteria, members of different genera in this family share less than 70% identity in their complete RdRP amino acid sequences (Ayllón et al., 2020). Based on the annotation, the Myrmica rubra ourmia-like virus 1 RdRp protein sequence shared 38% identical amino acids with the closest virus in the database, which suggests that at least a new genus could be formed.

### Pisuviricota

Pisuviricota is a phylum of positive single-stranded and double-stranded RNA viruses (Koonin et al. 2024). Pisuviricota was the overwhelmingly largest group in this study, and it is known that especially picornaviruses are very common in ants and Hymenoptera in general (Viljakainen et al. 2023). We show two examples for this group — one from the order Picornavirales and one outside of it. In the group outside of Picornavirales, there were 29 viruses (Supplementary Figure 2), of which the following representative is an example:

Temnothorax nylanderi partiti-like virus 1. The phylogenetic tree places this virus closest to the Partitiviridae family. These viruses infect plants, fungi, and protozoa. They are double-stranded RNA viruses with a 3-4.8 kb long bisegmented genome, where one segment contains the RdRp gene and the other has the capsid protein (Vainio et al., 2018). Temnothorax nylanderi partiti-like virus 1 has a 2111-nucleotide-long segment containing the RdRp gene, which falls into the range of known genera (Figure 2). The other segment containing the capsid protein presumably exists among the RNA-seq data but was excluded from the process at the latest during the annotation process, since only those sequences with RdRp were included in the phylogenetic analysis. According to the demarcation criteria, viruses of the same genus share more than 24% RdRp identity, while identity within a species must be over 90% (Vainio et al., 2018). The annotation result gave the identity of 57%, so it presumably belongs to the same genus as the closest match, Wuhan cricket virus 2, but that virus still remains unclassified, so the genus remains uncertain for Temnothorax nylanderi partiti-like virus 1.

A notable group in this phylum was the one containing 17 virus sequences that were named as sobeli-like viruses. These viruses formed a monophyletic group that was closest to but separate from the control viruses in the order Sobelivirales with families Alvernaviridae, Solemoviridae, and Barnaviridae.

For the order Picornavirales, there were 121 viruses to classify (Supplementary Figure 3). We chose an already known virus as a representative: Lasius niger virus 1, which was also found in *L. niger.* Lasius niger virus 1 belongs to the family Polycipiviridae, which are known to infect ants and other arthropods (Olendraite et al., 2019). The genome is 10-12 kb long, positive-sense nonsegmented RNA. The genome contains 4 or more ORFs (Olendraite et al., 2019). Lasius niger virus 1 belongs to the genus *Sopolycivirus,* which has 5 ORFs and a small overlapping ORF2b (Olendraite et al., 2019). The genome found in this study was 12,071 nt long (Figure 2), while the Genbank genome (acc: MF041812.1) is 11,092 nt long. ORF finder did not find an obvious ORF2b, but the remaining five ORFs had an identity above 98%, while the demarcation criteria required at least 90%.

### Kitrinoviricota

Kitrinoviricota is a phylum of positive single-stranded RNA viruses (Koonin et al. 2024). In Kitrinoviricota, we detected 19 viruses (Supplementary Figure 4), of which we chose Messor barbarus tobamo-like virus 1 as an example. This virus belongs to the family Virgaviridae, which has a 6,3-13 kb long positive-sense RNA genome, monopartite in the genus *Tobamovirus* and multipartite in other genera. They are plant viruses that are known to spread mechanically or via protista or nematode vectors. The genome structure of Messor barbarus tobamo-like virus 1 suggests it belongs to the genus *Tobamovirus*. The monopartite genome contains four ORFs: the first and second ORFs form the replication proteins, the third is the movement protein, and the fourth is the capsid protein (Adams et al., 2017). The found genome is 6306 nt long, which falls in the range of 6,3-6,6 kb found in other tobamoviruses (Figure 2). The species demarcation criteria requires the whole genome nucleotide sequence to be less than 90% identical to the closest match to be considered a new species. In this case, the identity to the closest species was 88%, so it can be classified as a new species.

A notable family in this phylum was Nodaviridae, in which the genome is bipartite and the RdRp and capsid proteins are in different segments. The species demarcation is done by comparing the sequence identity between the capsid proteins (Sahul Hameed et al., 2019). In this study, only the segments harboring the RdRp gene were collected, so the segments containing the capsid genes are presumably among the RNA-seq data, but connecting the RdRps to the missing capsids was beyond the scope of this study. Within the six ant species that had Noda-like RdRp genes, three also had Noda-like capsid genes among the annotated segments. Five sequences fell to the family Nodaviridae, and their species status was estimated according to the RdRp similarities, which were at most 50%, so they were classified as new species.

### Duplornaviricota

Duplornaviricota is a phylum of double-stranded RNA viruses (Koonin et al. 2024). In this phylum, we found 10 viruses (Supplementary Figure 5), of which we chose Temnothorax nylanderi leboti-like virus 1 as an example. This virus belongs to the family Lebotiviridae, which has a 6.7-8.1 kbp long double-stranded RNA genome that contains two slightly overlapping ORFs. The first ORF codes for the capsid protein, and the latter codes for the RdRp. The viruses in this family infect invertebrates (Hughes, 2024). In Temnothorax nylanderi leboti-like virus 1, the genome is 6761 bp long and the two ORFs overlap (Figure 2). According to the demarcation criteria, viruses that share more than 70% RdRp identity are considered to be the same species (Hughes, 2024). In this annotation, the identity percent was 57%, making it a new species.

### Negarnaviricota

Negarnaviricota is a phylum of negative-sense single-stranded RNA viruses (Koonin et al. 2024). In this phylum, there were 12 ant viruses (Supplementary Figure 6). As an example, a known virus, Lasius neglectus virus 2, was chosen, which belongs to the family Rhabdoviridae and was found in the ant *L. niger*. This is a very large family with 56 genera and over 400 species. The viruses infect a wide range of vertebrates, invertebrates, and plants, and include well-known harmful viruses such as the rabies virus. The genome is a 9.8-16.1 kb long single-stranded negative-sense RNA that contains five ORFs (Walker et al. 2022). The Lasius neglectus virus 2 genome found in this study was 12,069 nt long and contains five ORFs (Figure 2). The virus belongs to the genus *Alphahymrhavirus,* whose demarcation criteria require the following: 10% aa sequence divergence in N protein, 10% aa sequence divergence in L protein, and 15% aa sequence divergence in G protein (Walker et al. 2022). The annotation results showed that every ORF had a maximum divergence of 2% (98% identity) to the closest match, which was Lasius neglectus virus 2, indicating that it is indeed the same species.

Interestingly, we noticed that the name “Lasius neglectus virus 2” is the common name of two different viruses. The virus *Alphahymrhavirus neglectus* is the one in the phylum Negarnaviricota and family Rhabdoviridae, while the other virus *Sopolycivirus betalasii* belongs to the phylum Pisuviricota and family Polycipiviridae. *S. betalasii* was discovered in 2016 while *A. neglectus* was discovered in 2020. Therefore, we propose that *S. betalasii* keeps the original name and *A. neglectus* gets a new common name: Lasius neglectus rhabdo-like virus 1.

### Identification of shared virus species across ant species

There were 6 separate reciprocal BLAST searches made for each group representing the five different virus phyla, and separately for the order Picornavirales. In total, there were 19 cases where virus sequences were combined into one species due to the sequence similarity, in other words, the same virus was found from several ant species or the ant had several variants of the same virus: Formica sobeli-like virus 1 was found from *F. aquilonia* and *F. pratensis*, Crematogaster levior sobeli-like virus 1 had two variants in *C. levior*, Pheidole megacephala solinvi-like virus 1 had two variants in *P. megacephala*, Myrmicinae polycipi-like virus 1 was found from *My. ruginodis* and *P. fervida*, Myrmica rubra ifla-like virus 1 had two variants in *My. rubra*, Myrmica ifla-like virus 2 had in total five variants; one in *My. ruginodis* and four in *My. scabrinodis*, Myrmica ifla-like virus 1 had four variants; one in *My. rubra*, two in *My. ruginodis* and one in *My. scabrinodis*, Crematogaster levior picorna-like virus 1 had six variants in *C. levior*, known virus Hubei picorna-like virus 15, was found from *My. rubra* and *L. niger*, Myrmicinae picorna-like virus 1 was found from *Me. barbarus*, *My. rubra*, *My. ruginodis*, and *My. scabrinodis*, Myrmicinae picorna-like virus 2 was found from *Tet. bicarinatum* and *P. megacephala*, Myrmicinae dicistro-like virus 1 was found from *Me. structor* and *Tet. caespitum*, Myrmica dicistro-like virus 1 was found from *My. rubra* and *My. scabrinodis*, Myrmica picorna-like virus 1 was found from *My. rubra* and *My. ruginodis*, Myrmicinae noda-like virus 1 was found from *Me. structor* and *My. rubra*, Myrmicinae virus 1 was found from *Me. structor* and *P. megacephala*, Formica spinareo-like virus 1 was found from *F. aquilonia* and *F. pratensis*, known virus Chiqui virus was found from *Me. structor* and *My. scabrinodis*, and lastly Formica xinmo-like virus 1 was found from *F. aquilonia* and *F. pratensis*.

After combining the virus species, when applicable, the numbers in virus names were readjusted afterwards to keep the numerical order consistent. For example, after Formica pratensis sobeli-like virus 1 and Formica aquilonia sobeli-like virus 1 were combined into Formica sobeli-like virus 1, the next virus in the numerical order, Formica pratensis sobeli-like virus 2, was renumbered as 1 to keep the numerical order consistent. In total, 12 viruses had their numerical order readjusted (Table 5). Known viruses were classified as variants of a given virus if the whole genome nucleotide identity was below 95%, unless an official criterion for variant classification was available. Novel viruses grouped into a single species were assigned numbered variant names to distinguish individual genomes when submitted to GenBank.

**Table 5.**
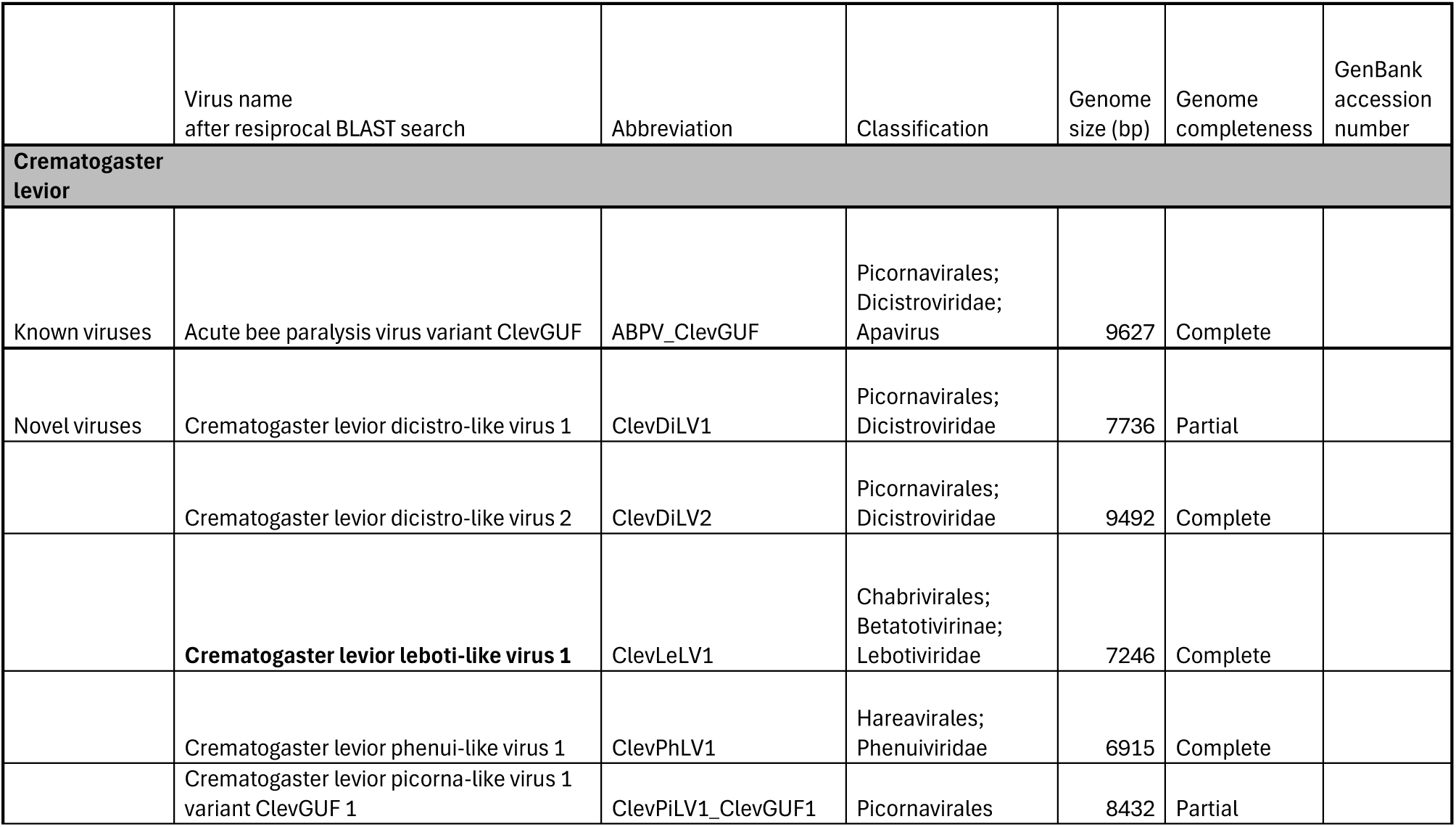
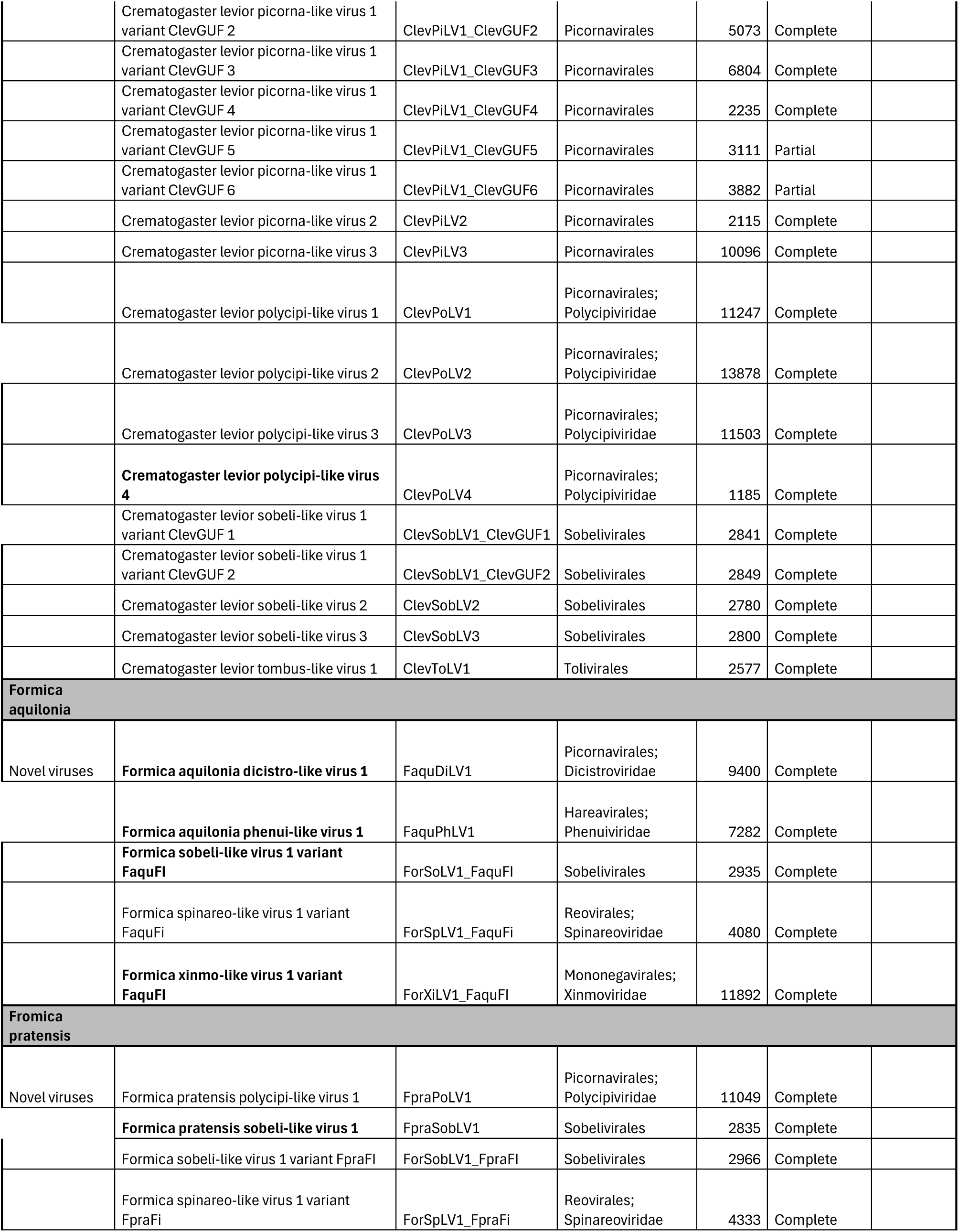
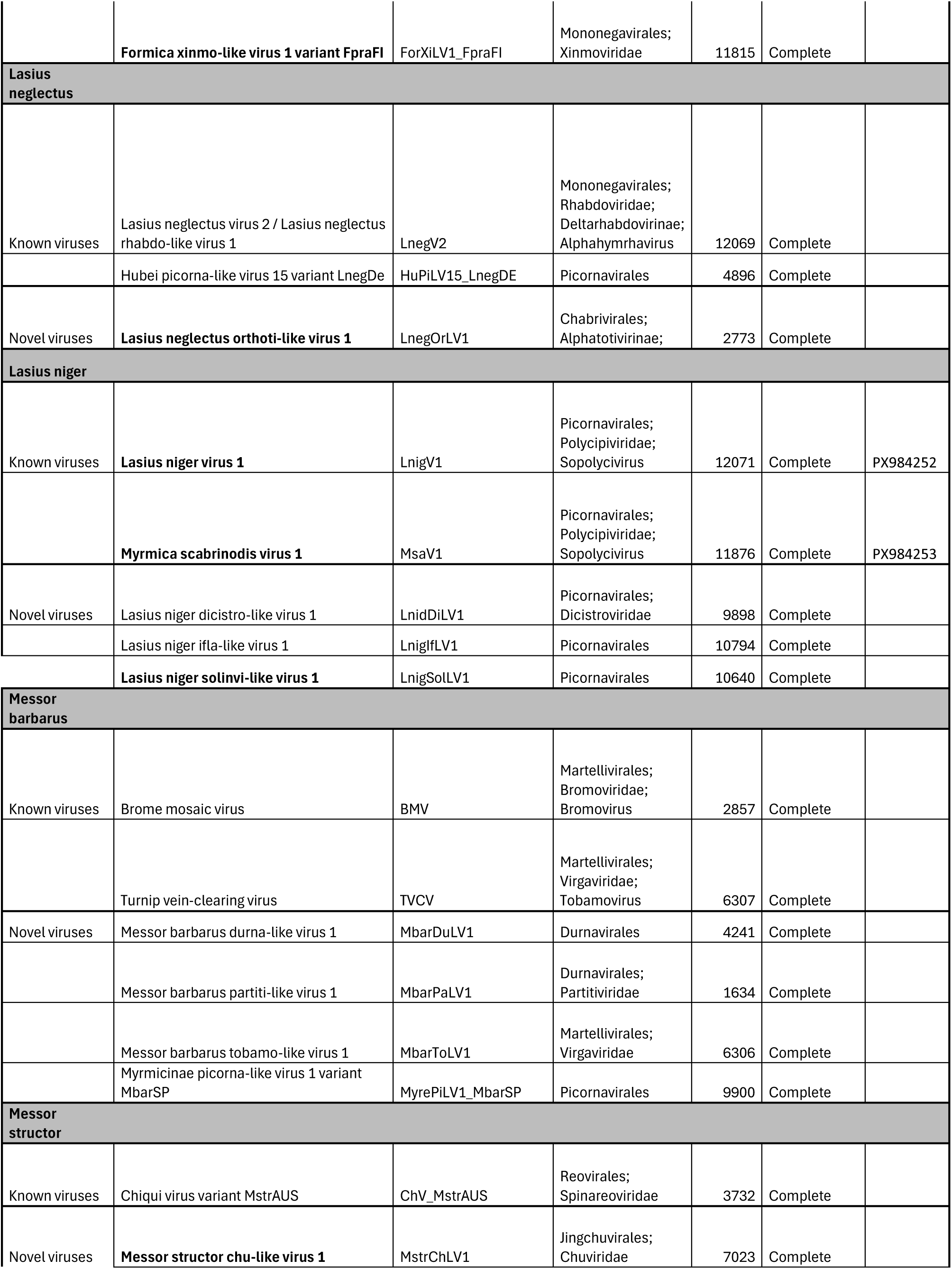
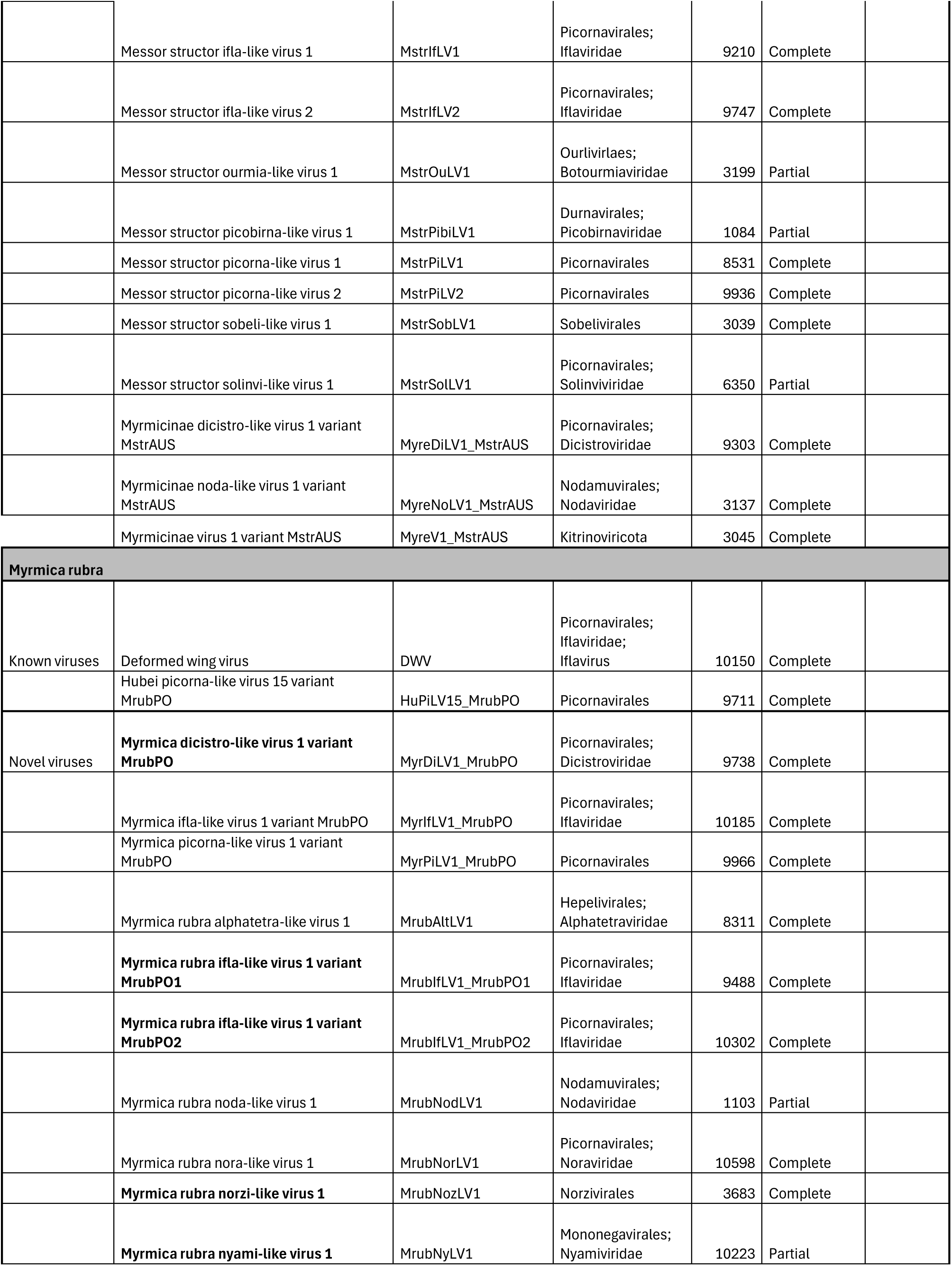
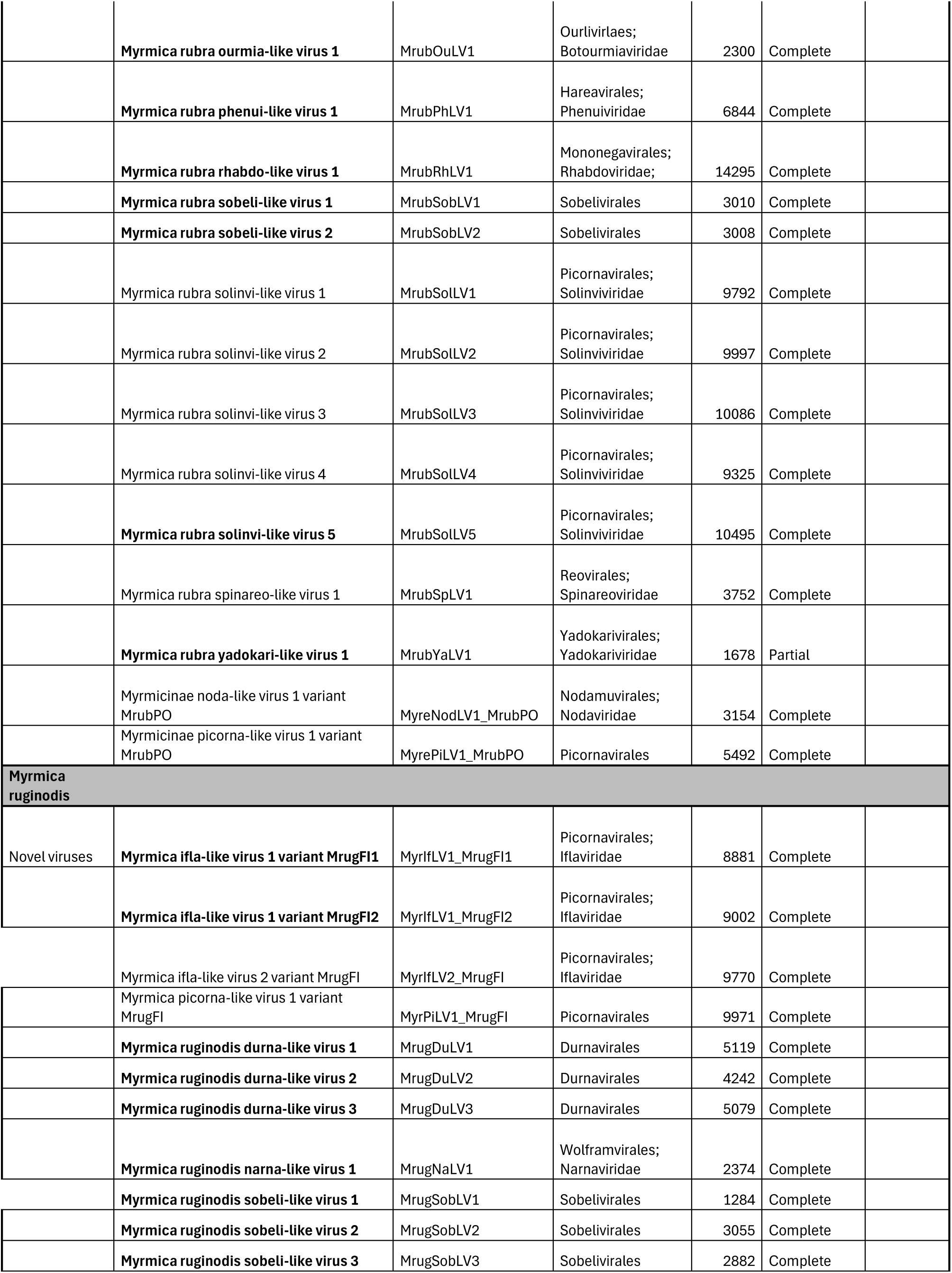
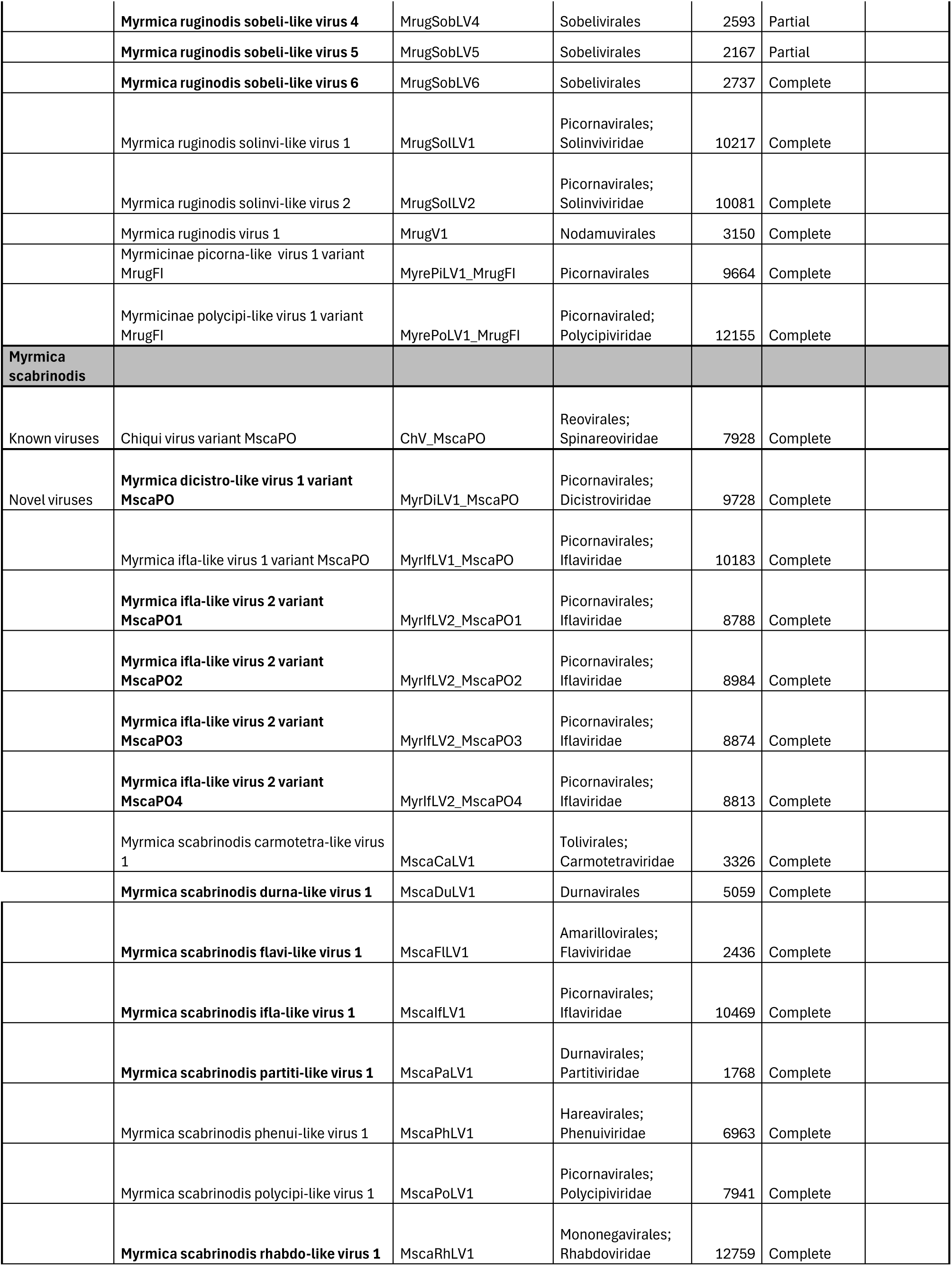
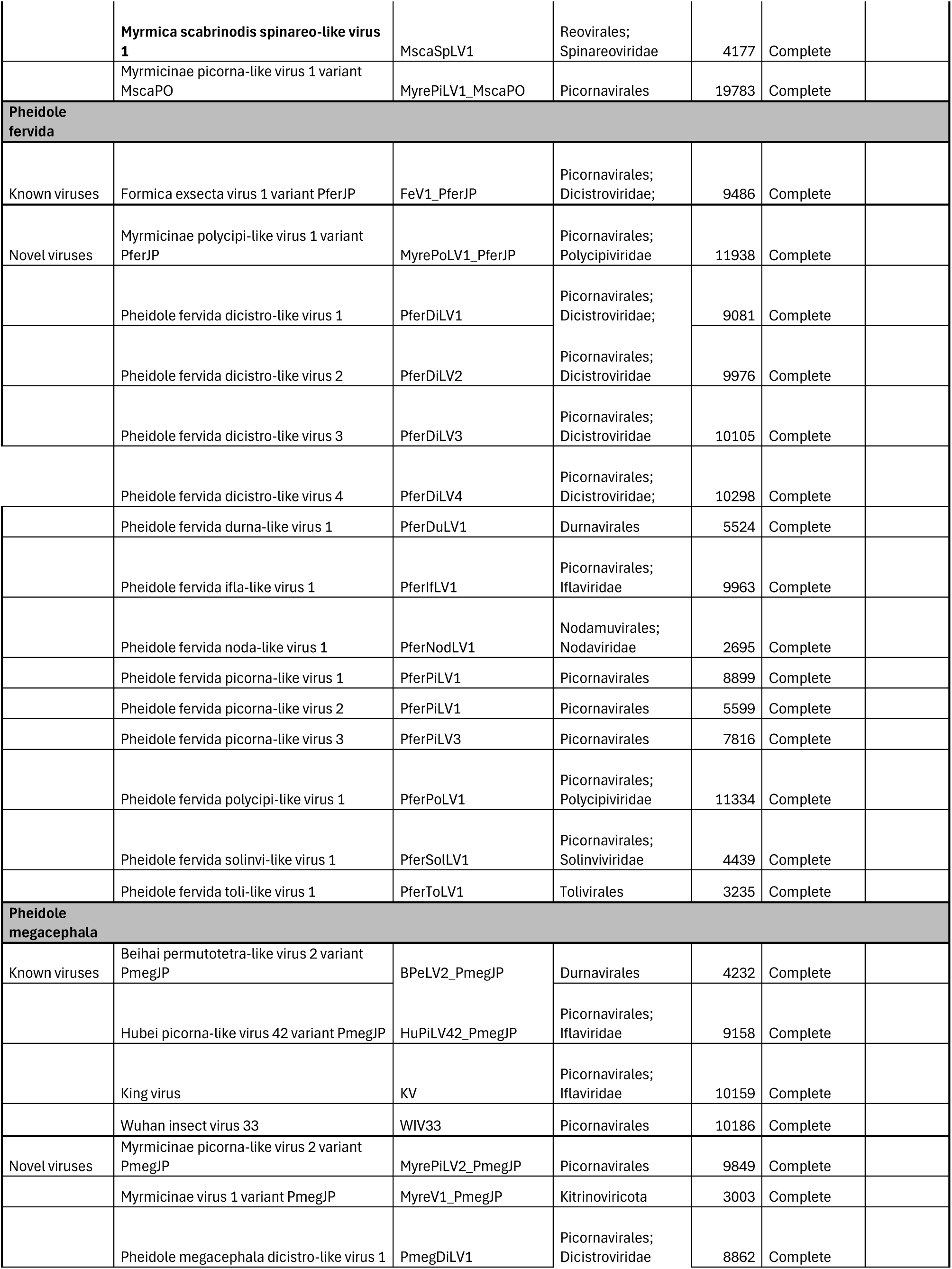
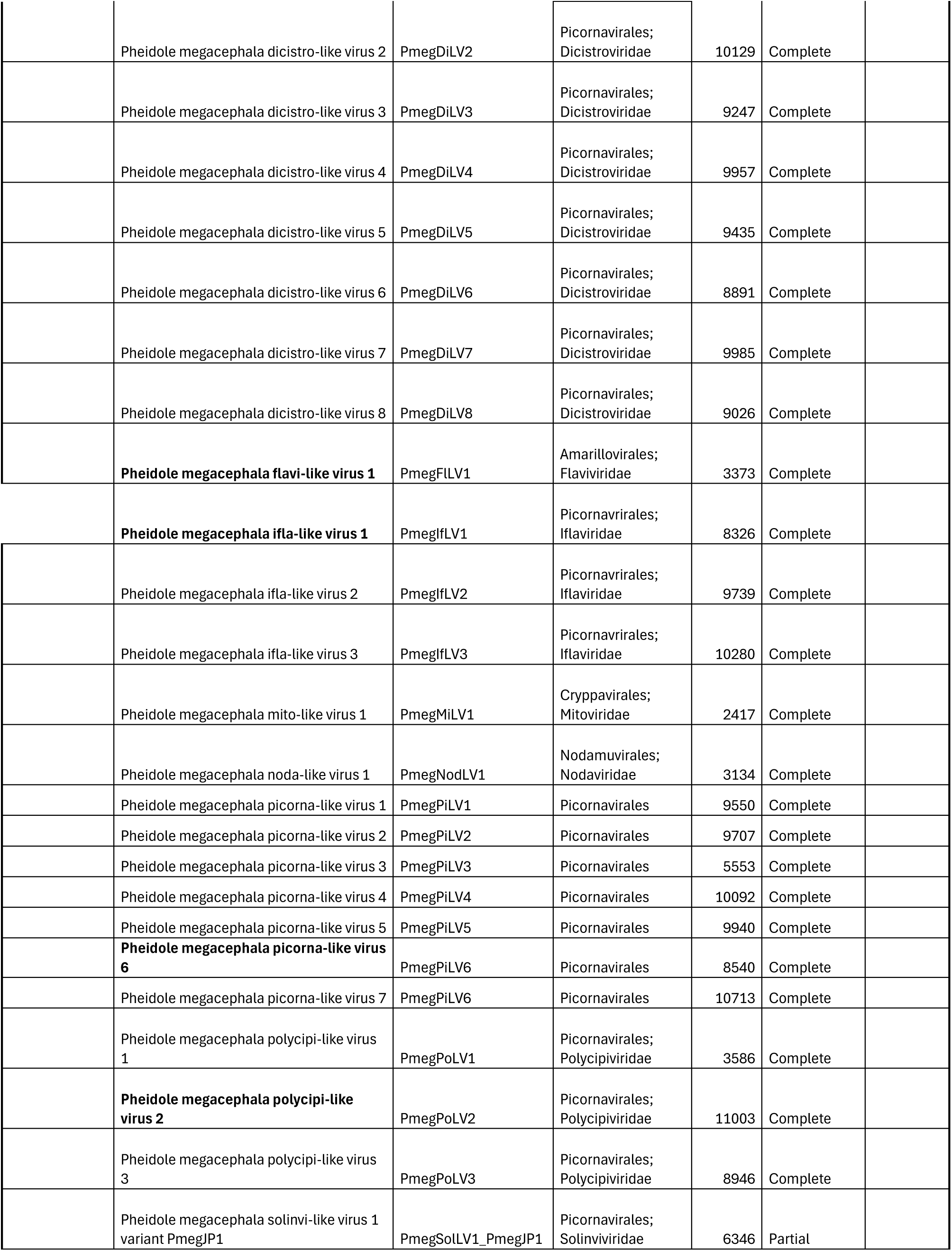
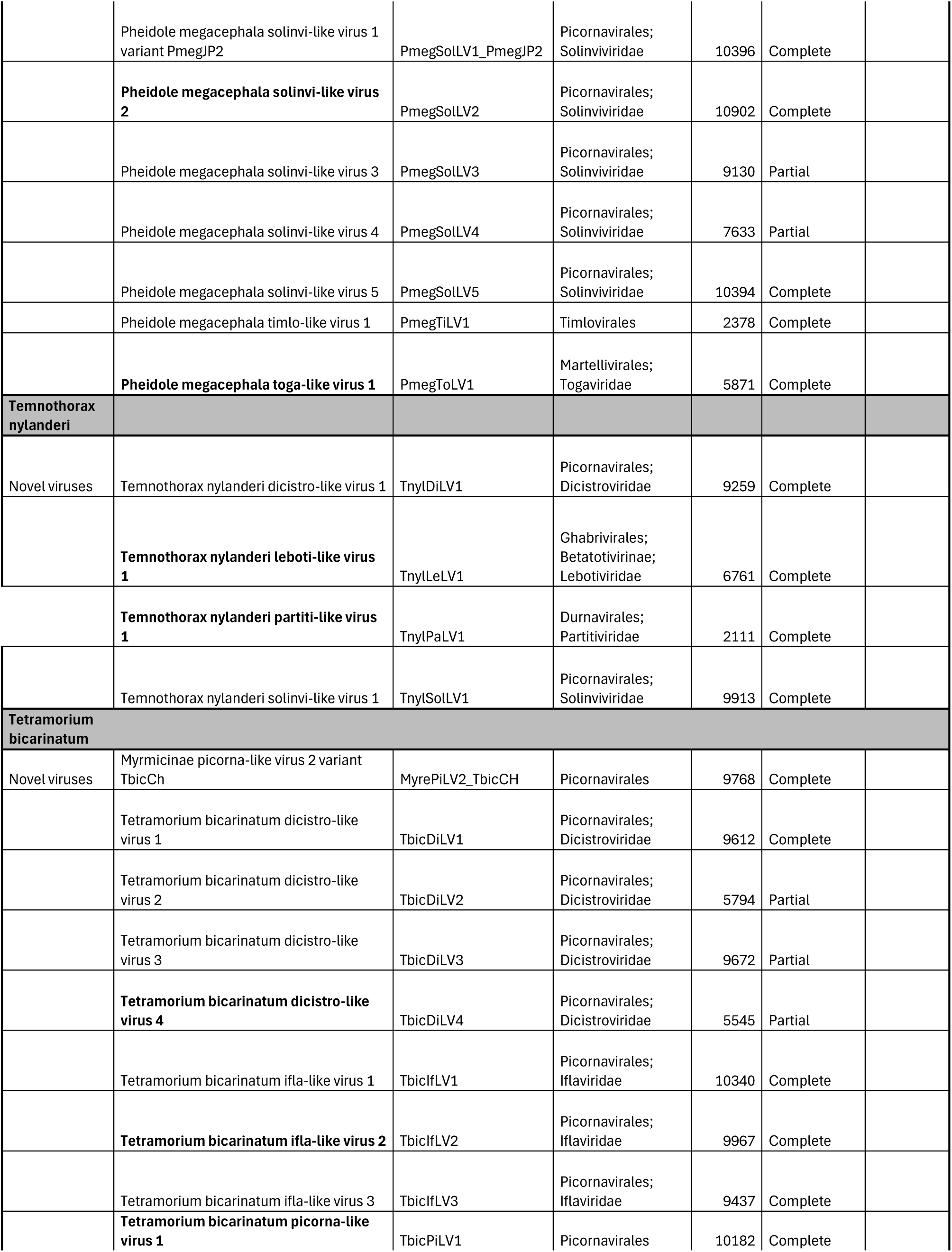
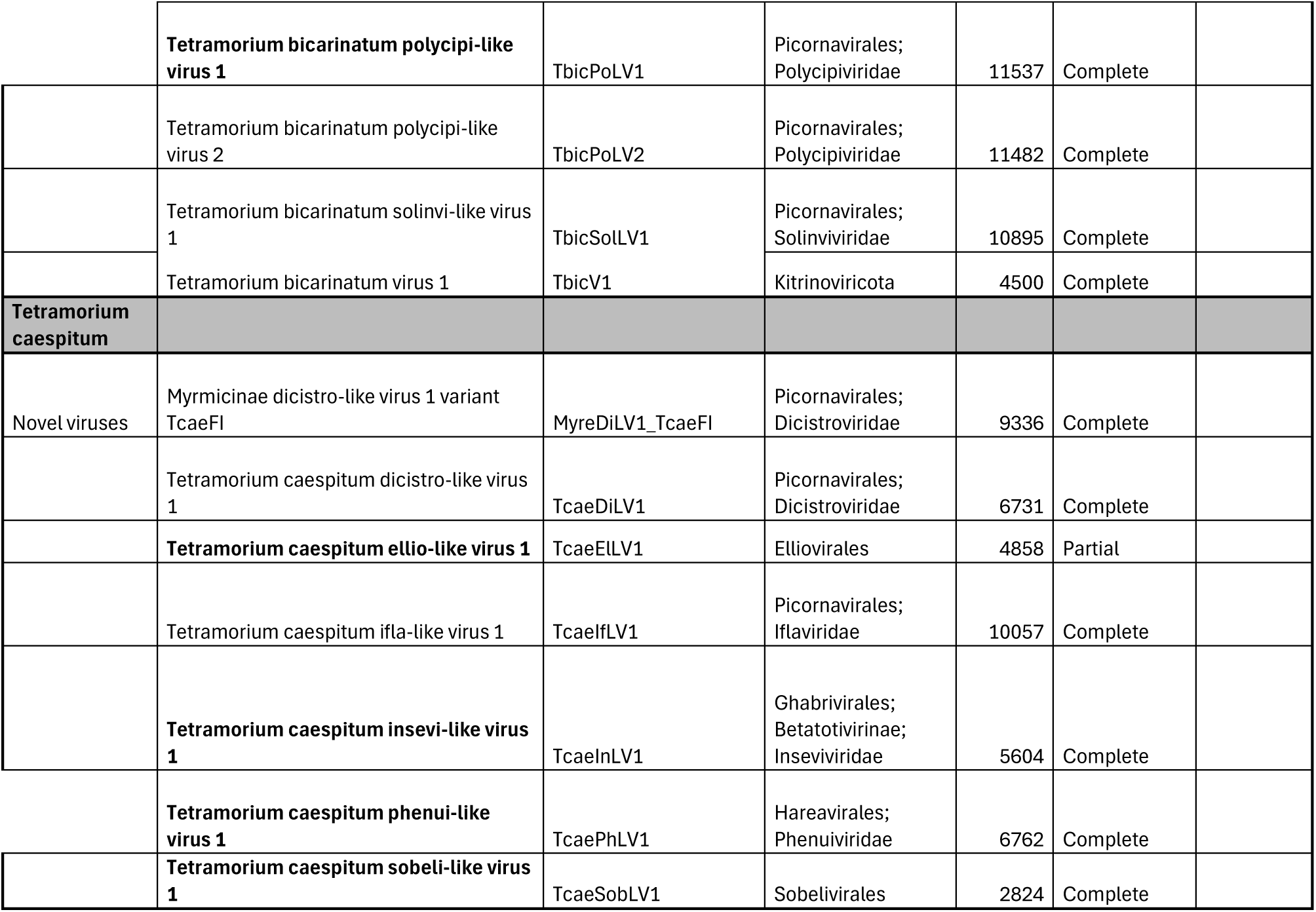
Known and novel viruses found in this study. Viruses in bold exhibited a positive RNAi sponse i.e. their siRNA RPKM value was above 10. Known viruses with whole genome identity less than 95% were marked as variants unless an official criteria for variants was available. Novel viruses grouped into a single species were named as variants to distinguish individual genomes when submitted to GenBank. Classification is provided to the closest taxonomix rank available.

This combination of species reduced the final number of viruses from 198 to 168, of which 154 (91.7%) were new species, and 14 (8.3%) were known viruses. Figure 3 shows the distribution of different viral phyla across ant species.

**Figure 3.**
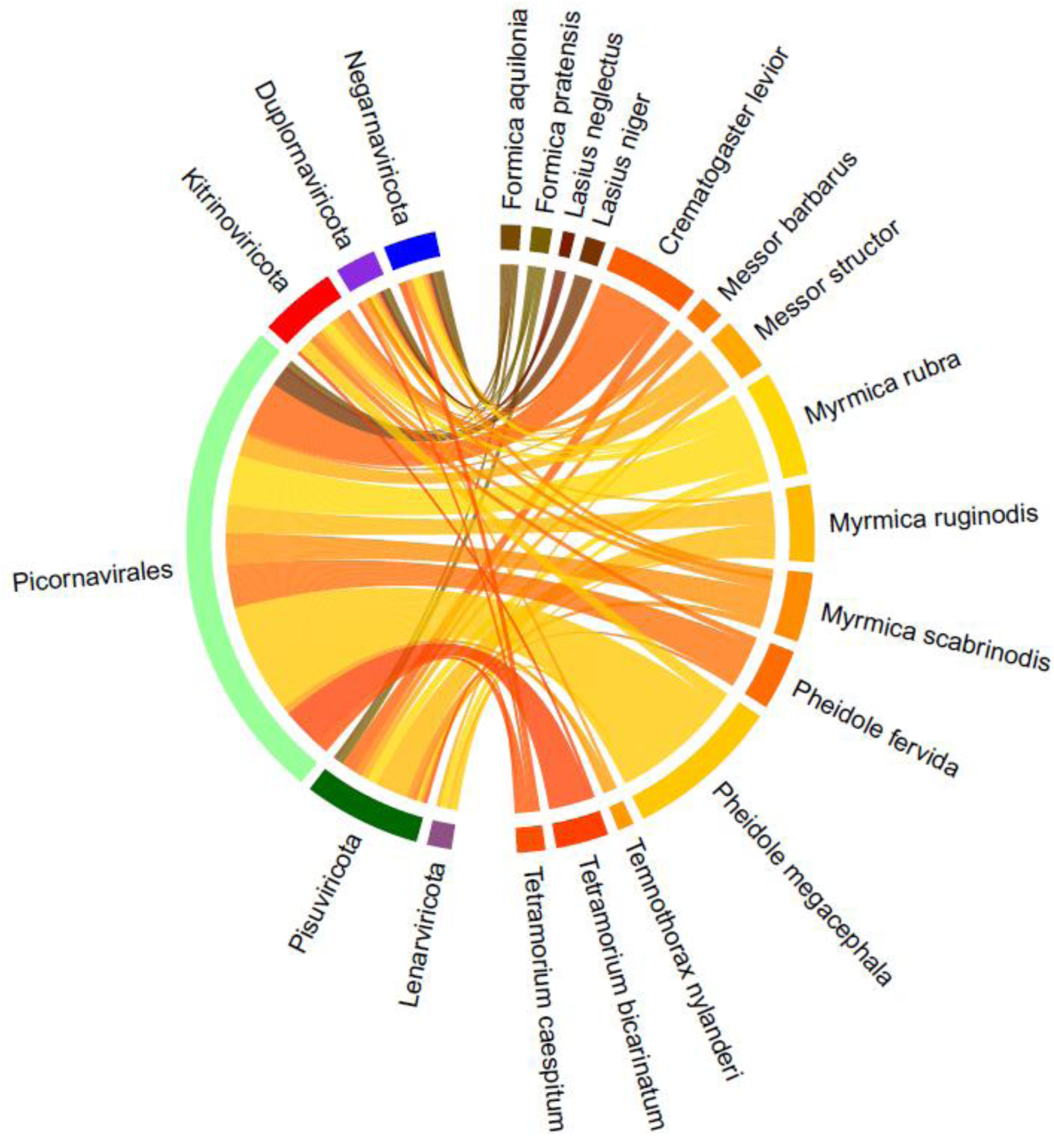
The distribution of viruses in ants. On the left side of the graph, there are the five virus phyla plus the order Picornavirales (light green) separated as its own group. On the right side of the graph are the fifteen ant species. The yellow to orange shades are ant species belonging to the subfamily Myrmicinae, and the ants in brownish shades belong to the subfamily Formicinae.

### RNAi-based immune response against the viruses

The sRNA size distribution was used to detect viruses that triggered an RNAi response in the host, which was indicated by a distinct peak of 21–22 nt sRNAs. Among the 168 virus species that were taxonomically classified, 59 were actively replicating based on the RNAi response.

The sRNA size distribution became increasingly random as the sRNA RPKM value decreased, even when the long RNA RPKM remained high. The relatively low number of actively replicating viruses implies contamination (from food or environment) or that some viruses might be highly abundant and actively replicating in the ant host, but they do not induce an RNAi response. Such viruses may therefore be able to evade recognition by the host RNAi machinery.

Of the remaining 97 viruses lacking an identified RdRp, 45 had a visible 21-22 nt peak. These viruses represent either partial virus genomes where the RdRp region is missing or viruses with segmented genomes where the RdRp is located in another segment.

### Identification of viruses based on the sRNA data

In addition to the 150 bp long RNA-sequences, viruses were separately assembled from the small RNA sequence data and aligned against a reference database of known viruses (GenBank gbvrl) using VirusDetect. The software provided the sequences of the aligned known viruses (either a whole genome, partial genome, or separate genes that are available in the reference database) and a visual representation of how the sRNA contigs aligned with the known sequences, i.e., how long the sRNA contigs are and what part of the known virus genome they match with (Figure 4).

**Figure 4:**
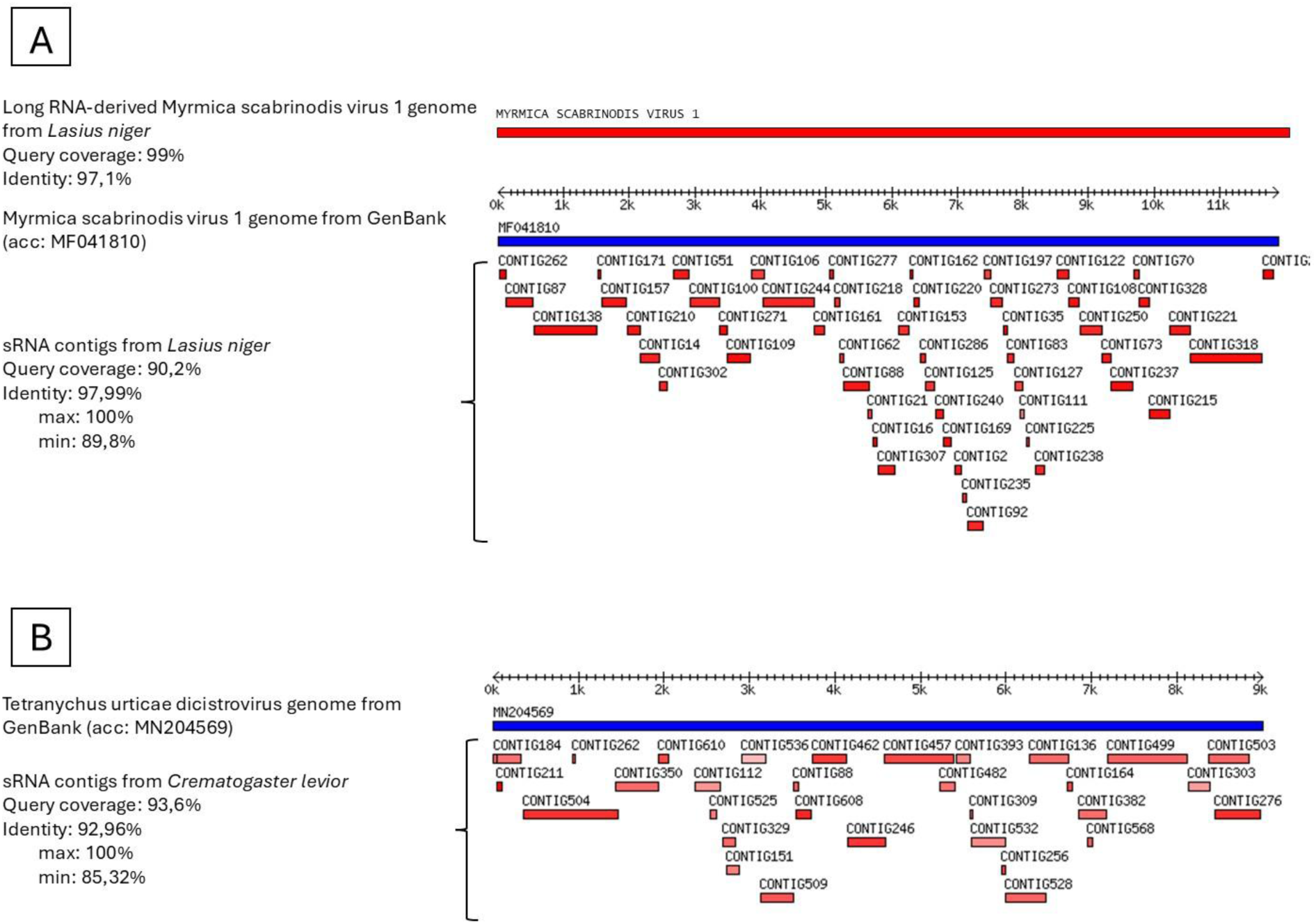
Two examples of VirusDetect results. A) VirusDetect visualization of *L. niger* sRNA contigs aligned against the known virus Myrmica scabrinodis virus 1 (acc: MF041810) with coverage of 90.2% and average identity of 97.99%. The above alignment is a manually added visualization of the long RNA-derived Myrmica scabrinodis virus 1 genome found from *L. niger* with coverage of 99% and identity of 97.1% to the known virus (MF041810). B) VirusDetect visualization of *C. levior* sRNA contigs aligned against the known virus Tetranychus urticae dicistrovirus with coverage of 93.6% and average identity of 92.96%. No matching viruses were found from long RNA-derived genomes; therefore, this virus can only be found from sRNA data.

Out of the 21 high-quality virus sequences used as a BLAST query against the longRNA-derived virus genomes, 9 matched the genomes from the same ant species. For example, from *L. niger*, VirusDetect found 51 sRNA contigs that covered 90.2% of a known virus’s, Myrmica scabrinodis virus 1, genome (GenBank accession number: MF041810) with 98% identity (min 89.8%, max 100%). When used as a query in the BLAST search, this virus (MF041810) matched the long RNA-derived virus Myrmica scabrinodis virus 1 genome with a coverage of 99% and an identity of 97%, indicating that the 51 sRNA contigs originated from the long RNA-derived Myrmica scabrinodis virus 1 (Figure 4A).

Out of the 21 high-quality virus sequences, 12 did not match with any long RNA-derived virus genome. These 12 sequences represented four different viruses found in the ants *C. levior, Me. barbarus,* and *Tet. caespitum*. These viruses were Tetranychus urticae dicistrovirus (Figure 4B), Tetranychus urticae-associated picorna-like virus 1, Tetranychus urticae-associated dicistrovirus 2, and Aphis glycines virus 1. *Tetranychus urticae* is a plant-feeding mite, and *Aphis glycines* is an aphid. Both mites and aphids are known to be associated with ants: mites as parasites (Campbell et al, 2013) and aphids in a mutualistic relationship (Stadler & Dixon, 2005). Close contact could offer a path for viruses to infect ants through these insects.

### Polygynous ants have higher virus diversity than monogynous ants

When comparing the monogynous and polygynous ant species, there were no statistically significant differences in the number of virus species between the groups (Wilcox rank sum test, W = 21, p-value = 0.2214), nor in total viral abundances, as approximated by RPKM (Wilcox rank sum test, W = 19, p-value = 0.1004). However, the visualized data suggest an overall trend: in both virus diversity (Figure 5A) and total viral abundance (Figure 5B), monogynous species generally appear to have fewer viruses than polygynous species, while the polymorphic species fall between these two groups. Even though the variation within each group remains substantial, the overall pattern suggests lower viral diversity and abundance in monogynous ants compared with polygynous ants.

**Figure 5.**
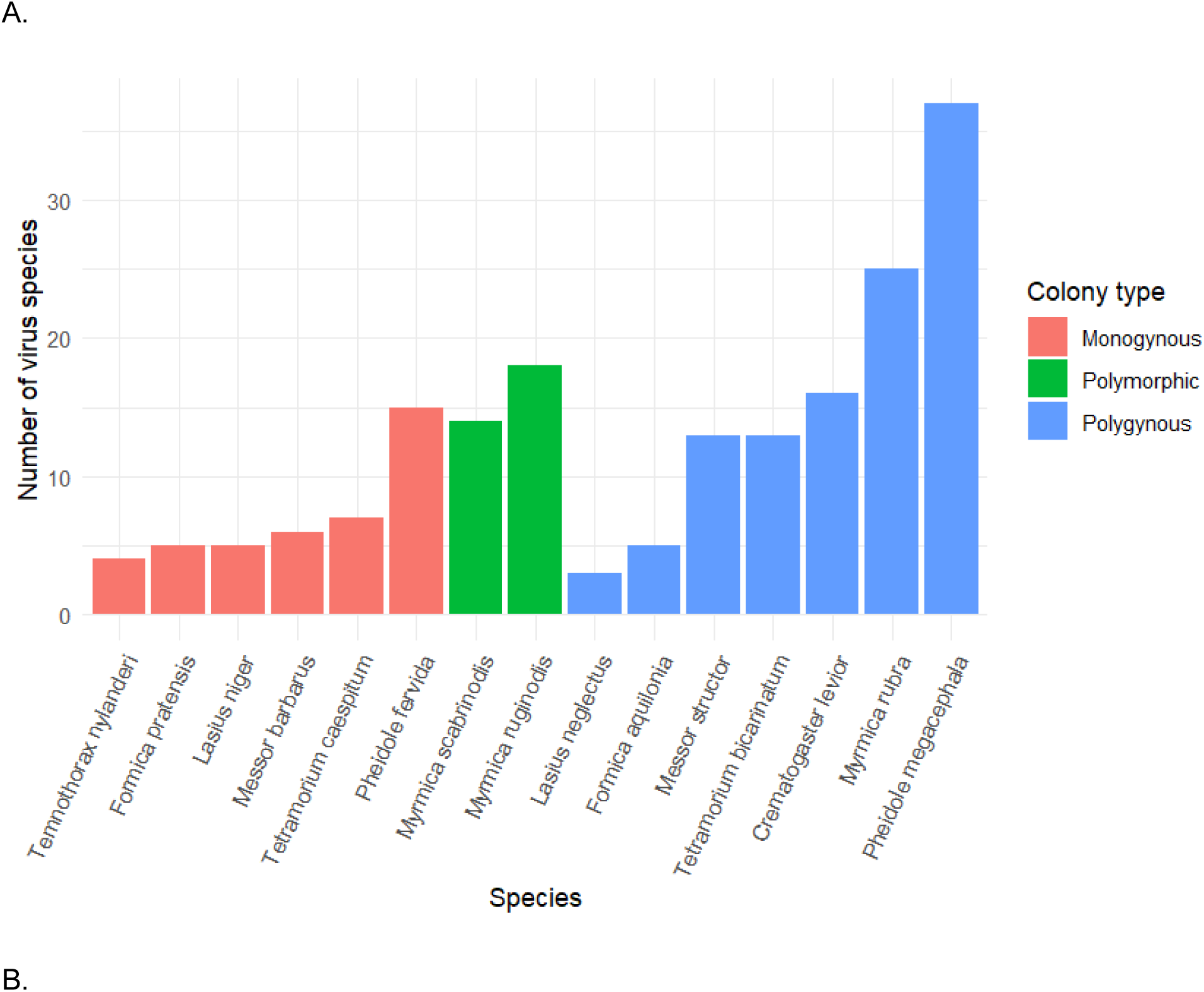
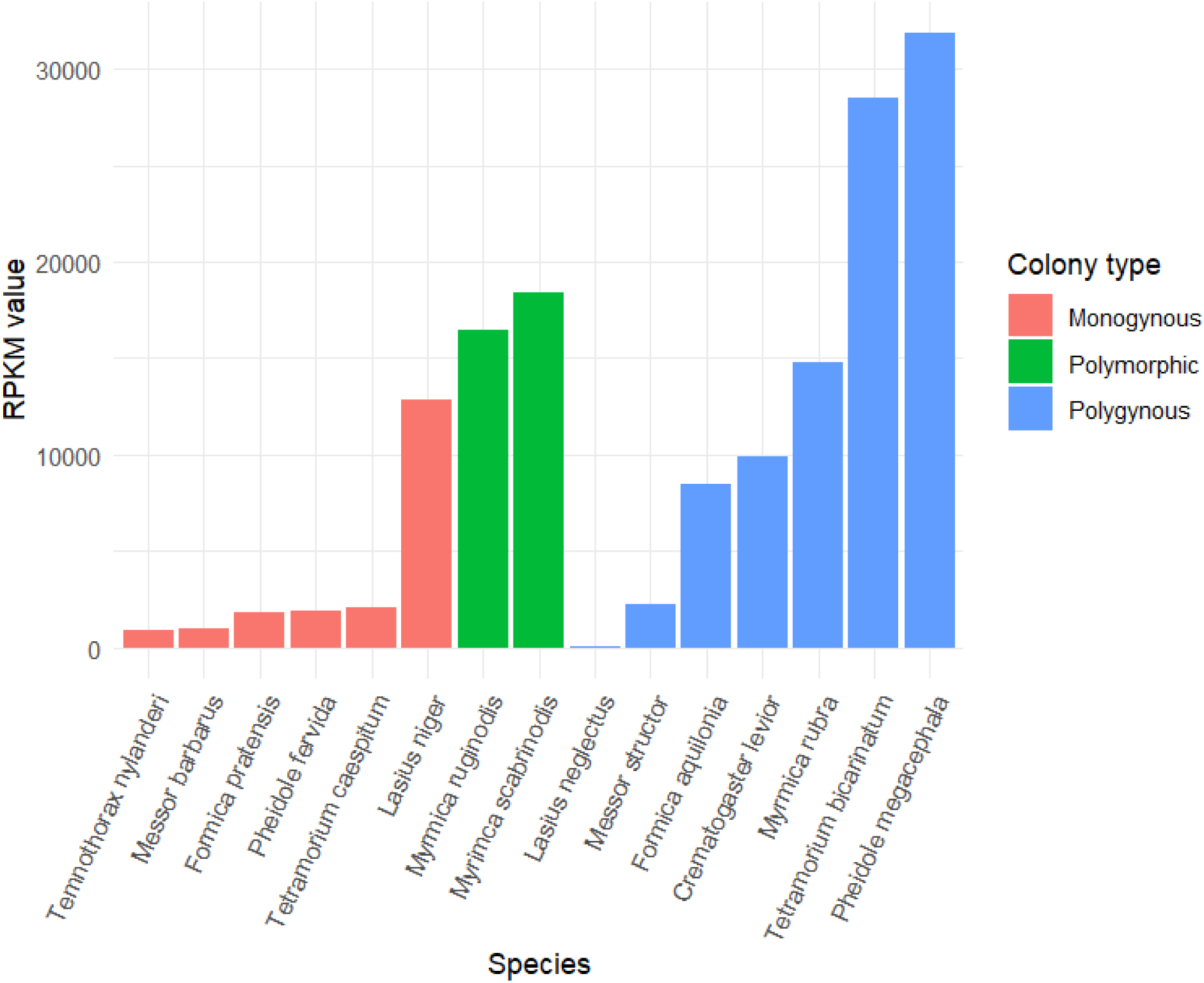
A. The number of virus species in each ant species. B. The total viral abundances in each ant species. The long RNA RPKM values of each individual virus in each ant species were summed together to get the total viral abundance.

When the ant species were divided into five closely related species pairs, the negative binomial generalized linear mixed model revealed that polygynous ants do have significantly more virus species compared to monogynous species (estimate = 0.63 ± 0.20 SE, z = 3.11, p = 0.0019). The estimated rate ratio indicated that polygynous species had, on average, 1.87 times more virus species than monogynous species (95% CI = 1.26 - 2.77). Species pair identity was included as a random effect to account for phylogenetic pairing. Model diagnostics using DHARMa indicated no evidence of overdispersion (dispersion test: p = 0.33). The paired Wilcoxon signed-rank test showed the same trend but was not statistically significant (V = 9, p = 0.20), likely due to the small number of species pairs (n = 5). Total viral abundance did not differ significantly between the two social structures. The linear mixed model on log-transformed viral abundance, with social structure as a fixed effect and species pair as a random effect, showed no significant difference between the groups (estimate = 0.57 ± 1.17 SE, t = 0.49). Similarly, the paired Wilcoxon signed-rank test did not detect a significant difference in the total viral abundance between monogynous and polygynous species (V = 12, p = 0.31).

When testing whether the higher number of virus species in the ants also correlated with overall higher viral abundances, a moderate positive correlation was detected, although the relationship was not statistically significant (Spearman’s ρ = 0.50, p = 0.14) (Figure 6).

**Figure 6:**
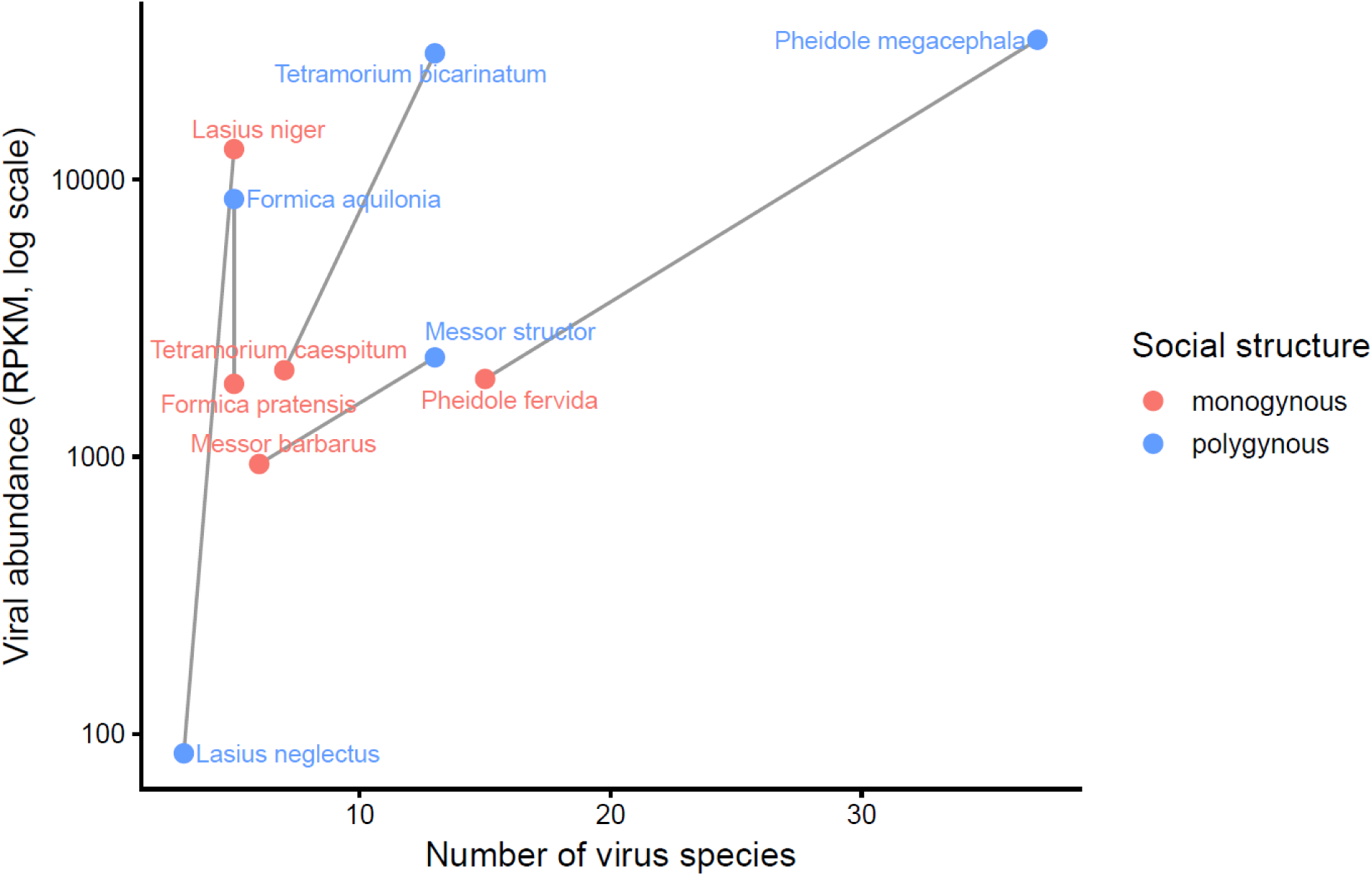
The correlation between the number of virus species and total viral abundance between the ant species pairs. The x-axis shows how many virus species were found in the ant, and the y-axis shows the combined RPKM value of these found viruses. The lines connect the monogynous (red) ant to their closely related polygynous (blue) species pairs.

When inspecting the viral abundances at the level of individual viruses (comparing a total of 110 RPKM values within the 5 ant species pairs), there were no significant differences between social structures. The linear mixed model on log-transformed RPKM values, with social structure as a fixed effect and species as a random factor, showed no effect of social structure on viral abundance (estimate = 0.11 ± 0.71 SE). On average, viruses were 1.11 times more abundant in polygynous species. This means that even though there seem to be more virus species in polygynous ants compared to monogynous ants, the virus abundances do not differ between the social structures. Figure 7 shows how both social structures have very abundant viruses as well as very rare viruses.

**Figure 7:**
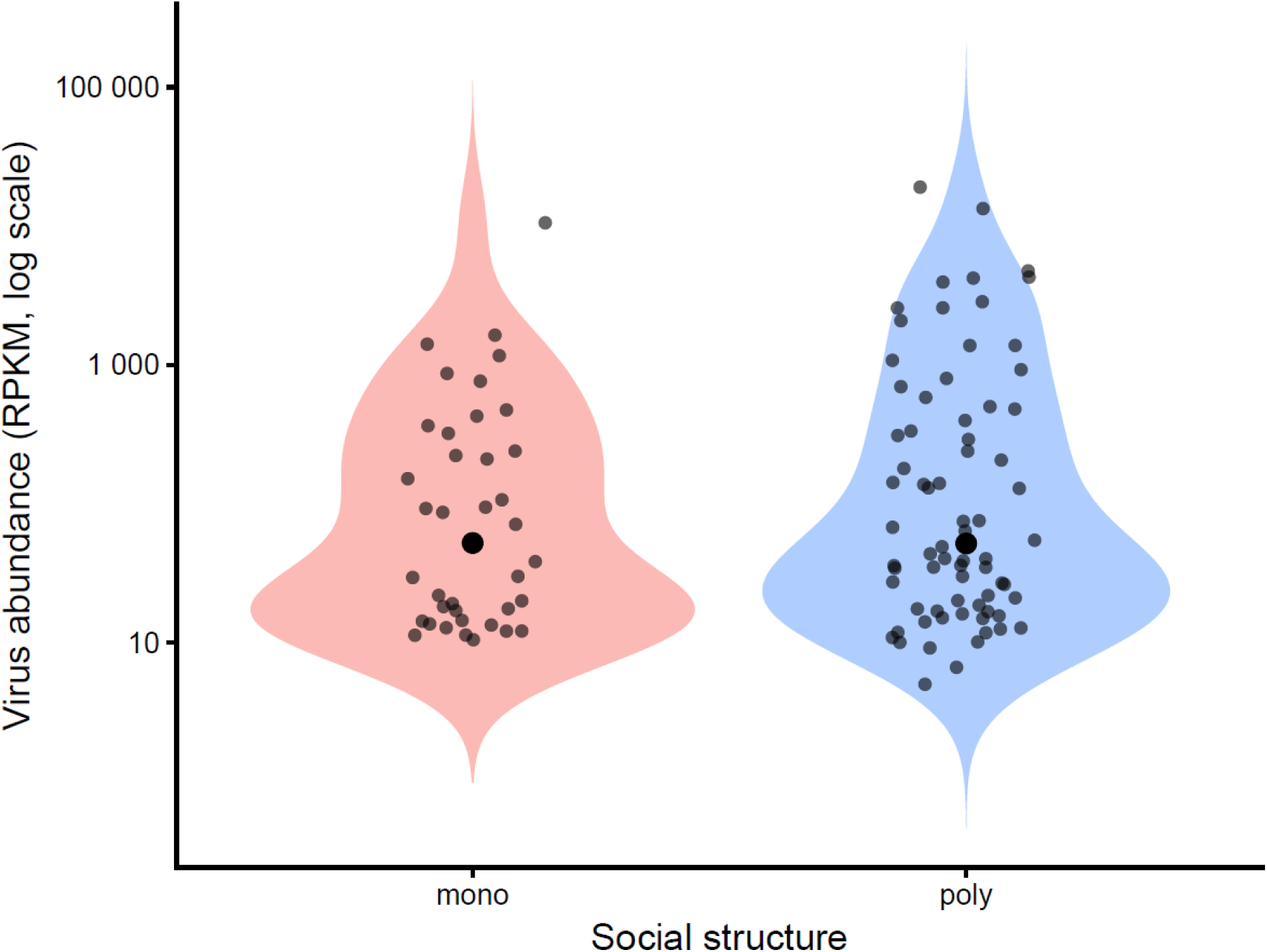
Virus abundance in RPKM values for individual viruses in monogynous and polygynous ant species. Both social structures have very abundant viruses and very rare viruses.

### Ant subfamily affects virus diversity

In our dataset of 15 ant species, Myrmicinae species harboured significantly more virus species than Formicinae species (Wilcox rank sum test, W = 3, P = 0.010). On average, Formicinae species contained 4.5 virus species, whereas Myrmicinae species contained 15.3 virus species (Figure 8). Viral community composition differed significantly between the two ant subfamilies (PERMANOVA, Bray–Curtis distance, F = 5.51, R² = 0.298, P = 0.013). Approximately 30% of the variation in viral species composition was explained by the ant subfamily. Pisuviricota viruses were significantly more diverse in Myrmicinae than in Formicinae (Wilcox rank sum test, W = 4, P = 0.016). NMDS ordination (stress = 0.072) showed a clear separation between Formicinae and Myrmicinae viral communities, consistent with the PERMANOVA results. Multivariate dispersion did not differ significantly between subfamilies (betadisper permutation test, P = 0.625), indicating that the PERMANOVA result reflected differences in viral community composition rather than unequal within-group variability.

**Figure 8.**
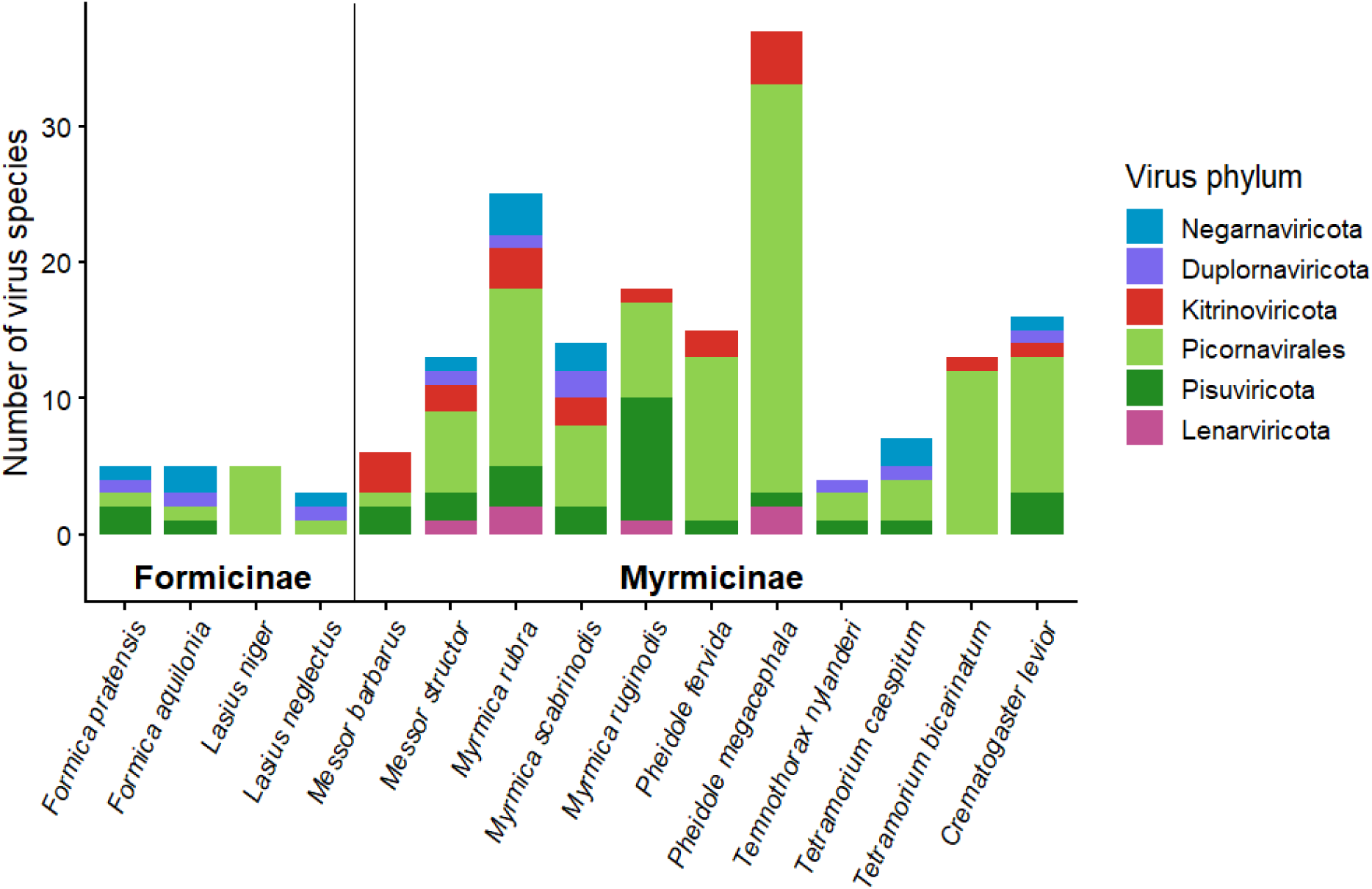
Virus phylum distribution between Formicinae and Myrmicinae. The bar height indicates the number of unique virus species found in each ant species, the color of the bar indicates the phylum in which the viruses belong to. Picornavirales (light green) is shown separately from the remainder of Pisuviricota (dark green) because it constitutes the most abundant viral order detected in the dataset.

## Discussion

In this study, we provide a comprehensive view of differences in RNA virus diversity, abundance, and infectivity in different types of ant societies. We characterized a total of 168 RNA viruses found in 15 ant species representing both monogynous and polygynous social structures. 35% (59) of these viruses caused an active infection, as determined by detecting the host ant RNAi-based immune response.

We were particularly interested in determining whether virus diversity and load differ between monogynous and polygynous ant species. We observed that polygynous ant species have a higher virus diversity than monogynous ant species. The result is consistent with the observation Brahma et al. (2022) made when comparing virus diversity among monogynous and polygynous *Solenopsis invicta* colonies. Brahma et al. (2022) also observed significantly higher viral abundances in polygynous colonies compared to monogynous colonies, whereas in our study, the viral abundances between polygynous and monogynous species did not differ. The discrepancy could be explained by our more diverse and varied dataset, which includes several ant species, whereas Brahma et al. studied colonies within a single species, where the high viral abundance could be a consistent trend. Differences in viral diversity between monogynous and polygynous species may be linked to their colony size. Polygynous colonies span larger areas and thus are exposed to more viruses. The lack of differences in viral abundances however implies that both social structures are similarly able to manage the virus infections.

Interestingly, the ant subfamily affects virus diversity. We observed that Myrmicinae ants (n=11) have more than three times more virus species (on average 15.3 species) compared to Formicinae ants (n=4) (4.5 species), this difference being statistically significant. The viral communities also differed between subfamilies, for example Lenarviricota and Kitrinoviricota viruses were only found in Myrmicinae ants. The absence of Lenarviricota and Kitrinoviricota in Formicinae ants could be explained by just the small number of viruses or small number of sampled ant species. When one finds fewer viruses, there is naturally less diversity. Or there could be something that makes Formicinae especially resistant to these phyla? A possible explanation for the overall lower number of viruses in Formicinae is the presence of an acidopore, a poison gland unique to this ant subfamily that produces formic acid (Ward, 2007, Koch et al., 2025). Formic acid can be sprayed as a defense against predators, and to disinfect the nest material, but it is also used to disinfect the brood and the ants themselves through a behavior known as acidopore grooming (Tragust et al., 2013, Koch et al., 2025).

As predicted, Picornavirales was the largest virus group in this study, the order alone containing 58% of all the found viruses. This observation confirms the earlier results of Viljakainen et al. (2023) and Zueva et al. (2025). There were several orders of viruses that have not been associated with ants before (Zueva et al., 2025, Flynn & Moreau, 2024): in Lenarviricota, there were orders Ourlivirales (previously associated with plants and fungi, 2 viruses) and Cryppavirales (fungi, 1 virus). In Pisuviricota, there were Yadokarivirales (fungi, 1 virus) and Sobelivirales (protista, fungi, and plants, 15 viruses). In Kitrinoviricota, there were Amarillovirales (mammals and birds with insect vector, 2 viruses). In Negarnaviricota, there were Elliovirales (crustacean, plants, mammals with insect vector, 1 virus). In these 22 viruses, a clear peak of 21-22 nt-long RNAs was visible in 18 viruses, suggesting that they might be infective. There was one virus, Formica xinmo-like virus 1, that can be classified into a new genus according to the demarcation criteria. In several cases, there were viruses that could very likely be classified into a new genus or even a new family judging by their sequence identity (Supplementary Table 1) or their placement in the phylogenetic trees (Supplementary Figures 1-6), but they did not have established genus demarcation criteria available.

Out of the 168 virus species found, 154 were new to science, adding to the list of almost 3900 known ant viruses (Zueva et al., 2025, Flynn & Moreau, 2024). Out of the 14 known viruses, 4 were viruses that have been first described in ants (Lasius niger virus 1, Myrmica scabrinodis virus 1, Formica exsecta virus 1, and Lasius neglectus virus 2 / Lasius neglectus rhabdo-like virus 1 (note the suggested new name). Out of the remaining 10 viruses, 8 were directly associated to wide range of Arthropoda, including bees (Beihai permutotetra-like virus 2, Deformed wing virus, King virus, Hubei picorna-like virus 42, Hubei picorna-like virus 15, Wuhan insect virus 33, Acute bee paralysis virus, and Chiqui virus), while the remaining two (Turnip vein-clearing virus and Brome mosaic virus) were mostly associated with plants but the listed sources/hosts in GenBank also included thrips, leafhoppers or “water from a ditch surrounding a field”. Most of the discovered viruses appear to be ant-specific, and most of those even ant species-specific, an observation Viljakainen et al. (2023) made as well. However, some of the viruses appeared to be able to infect a broader range of insects.

Viruses that were found in different ant species mostly originated from ants collected within the same country or directly neighboring countries or within Europe. However, there were two cases where the geographical distance was much wider: Myrmicinae polycipi-like virus 1 was found in *P. fervida* and *My. ruginodis,* which were collected from Japan and Finland, and Myrmicinae virus 1 was found in *Me. structor* and *P. megacephala,* which were collected from Austria and Japan. The fact that the same viruses can be found in ants that are either geographically or taxonomically so distant from one another warrants further research on the infection and transmission dynamics of the ant viruses.

When deciding which virus genomes should be selected for annotation, the RPKM value above 10 was used as a very strict criterion for determining what counts as a replicating virus. This way, the possibility of including contaminating viruses that are not replicating in the ant but originate from elsewhere, like the environment or the food the ant had eaten, could be mitigated, and the identified viruses would reliably be ant viruses. Especially in small RNA RPKM, the value of 10 proved to be an adequate threshold. When visualizing the size distribution of small RNAs with viRome, the clear sign of RNA interference seen in the abundance of 21-22 nucleotide-long sequences began to diminish very rapidly when the RPKM value went below 10, and the size distribution appeared either randomly scattered or nonexistent (Supplementary Table 1).

The large pool size of 400 individual ants per species might have led to a relatively abundant virus having an RPKM value below 10 and, therefore, being excluded from the analysis. For example, if only one of the 400 ants had a strong virus infection, the signal in the RPKM value might have been diluted so low that it’s no longer significant on a population scale, as prevalence within a colony can be variable (Okada et al., 2025, Manfredonia et al., 2026).Hence, this study highlights the most common ant viruses on a population scale. If the virus genomes with RPKM values below 10 were to be analysed, more rare viruses could potentially be found, though there would be an increasing likelihood of including contaminating viruses that would not be associated with ants in any way other than that they were found in the same place.

In conclusion, this study demonstrates that ants harbor a diverse range of viruses, most of which seem to be ant-specific viruses. As previously observed, Picornavirales is the most prevalent order of viruses associated with ants. We observed that polygynous ants harbor more virus species than monogynous ants. In the present dataset, the ant subfamily appeared to have a strong influence on virus diversity, with Formicinae harboring fewer viruses than Myrmicinae. This pattern may reflect the enhanced antimicrobial defenses of Formicinae, which use formic acid in colony hygiene and pathogen defense. Ultimately, our results reveal the impact of colony organization and lineage-specific traits on host-pathogen dynamics, paving the way for a deeper understanding of viral evolution in social insects.

## Supporting information

Supplementary Figure 1. Lenarviricota

Supplementary Figure 2. Pisuviricota

Supplementary Figure 3. Picornavirales

Supplementary Figure 4. Kitrinoviricota

Supplementary Figure 5. Duplornaviricota

Supplementary Figure 6. Negarnaviricota

Supplementary Table 1. Annotation

## Acknowledgments

We thank Soile Alatalo and Laura Törmälä for help with the laboratory work, Max Aubry, Iris Schlick-Steiner, Mihika Sen, Yue Shi, Yuya Suzuki, Paulina Chudzik, and Magdalena Witek for help with sample collection, Jasmin Oona Turunen for help with the data analysis and Elliot J. Lefkowitz for instructions on virus species classification.

## Funding

This work was supported by the Research Council of Finland grant no. 343022 to L.V. and Oulun Luonnonystäväin Yhdistys ry to M.Konu .

E.S. was supported by the Juan de la Cierva postdoctoral fellowship (grant no. JDC2024-054485-I) financed by the MICIU/AEI/10.13039/501100011033 and FSE+ J.O. was supported by an Investissement d’Avenir grant managed by the Agence Nationale de la Recherche (CEBA, ANR-10-LABX-25-01)

D.G. has benefited from the equipment and framework of the COMP-R Initiatives, funded by the “Departments of Excellence” program of the Italian Ministry for University and Research (MUR, 2023–2027).

S.L. acknowledges support from the Dutch Research Council (NWO; Grant OCENW. XS22.4.111).

## Notes

### Competing Interest Statement

The authors have declared no competing interest.

