## Supplementary figures and images for "Viral infection patterns in ants are affected by colony structure and phylogenetic lineage"

### Supplementary Figure 1. Lenarviricota

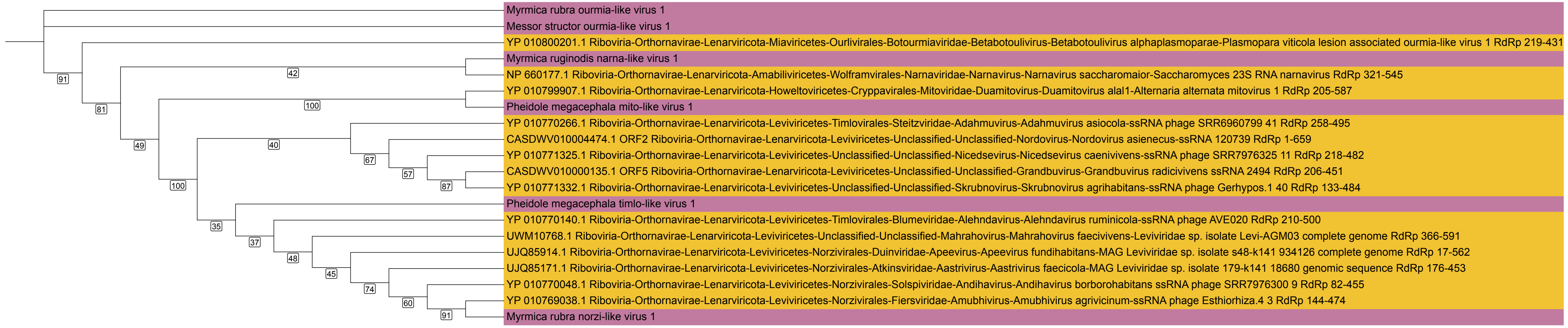

### Supplementary Figure 2. Pisuviricota

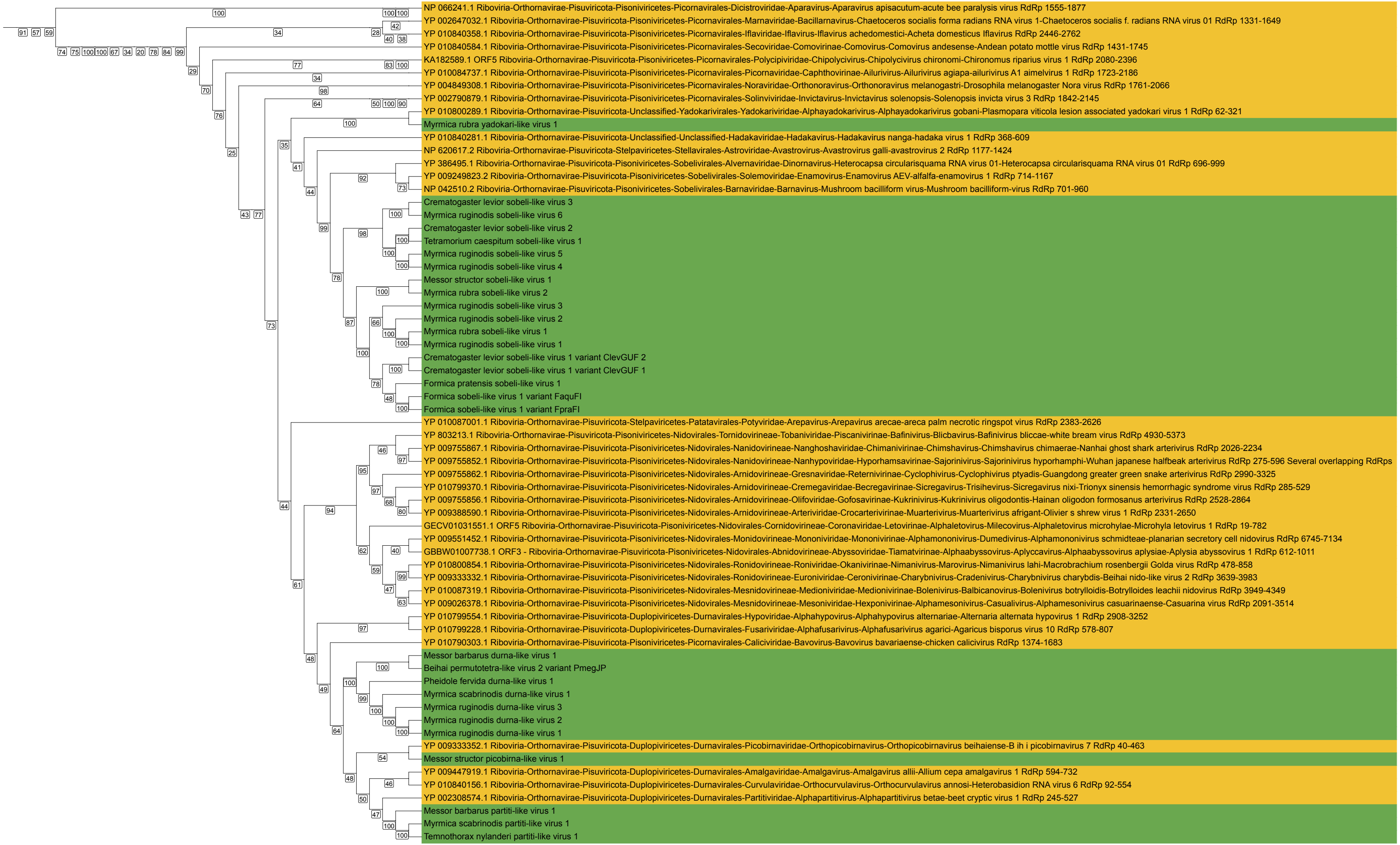

### Supplementary Figure 3. Picornavirales

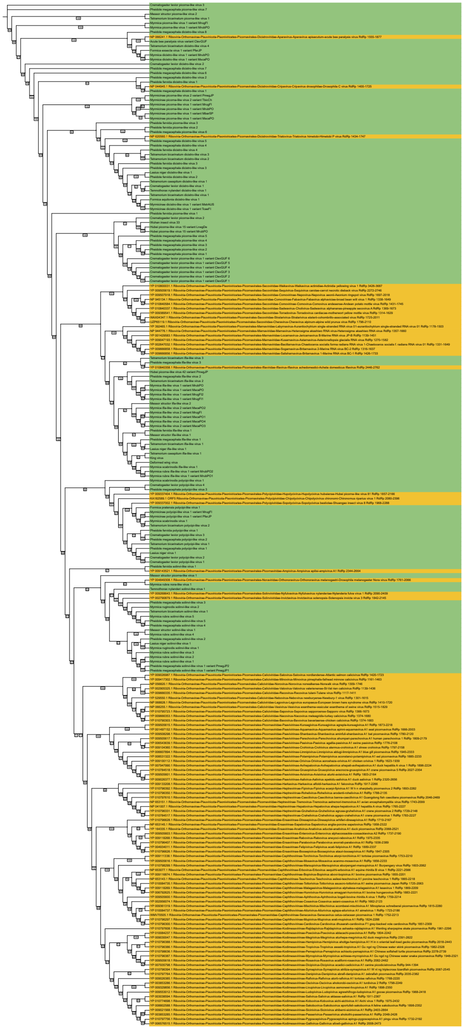

### Supplementary Figure 4. Kitrinoviricota

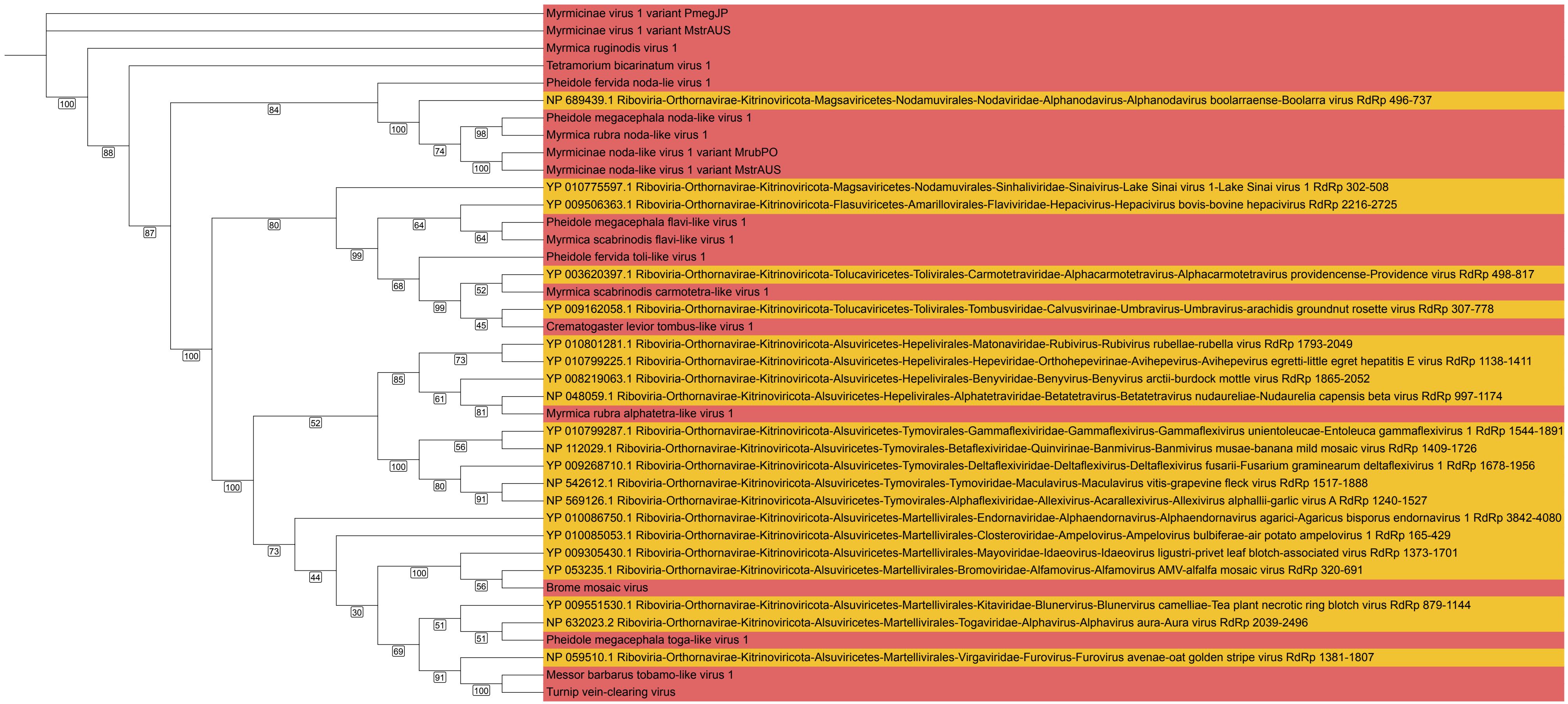

### Supplementary Figure 5. Duplornaviricota

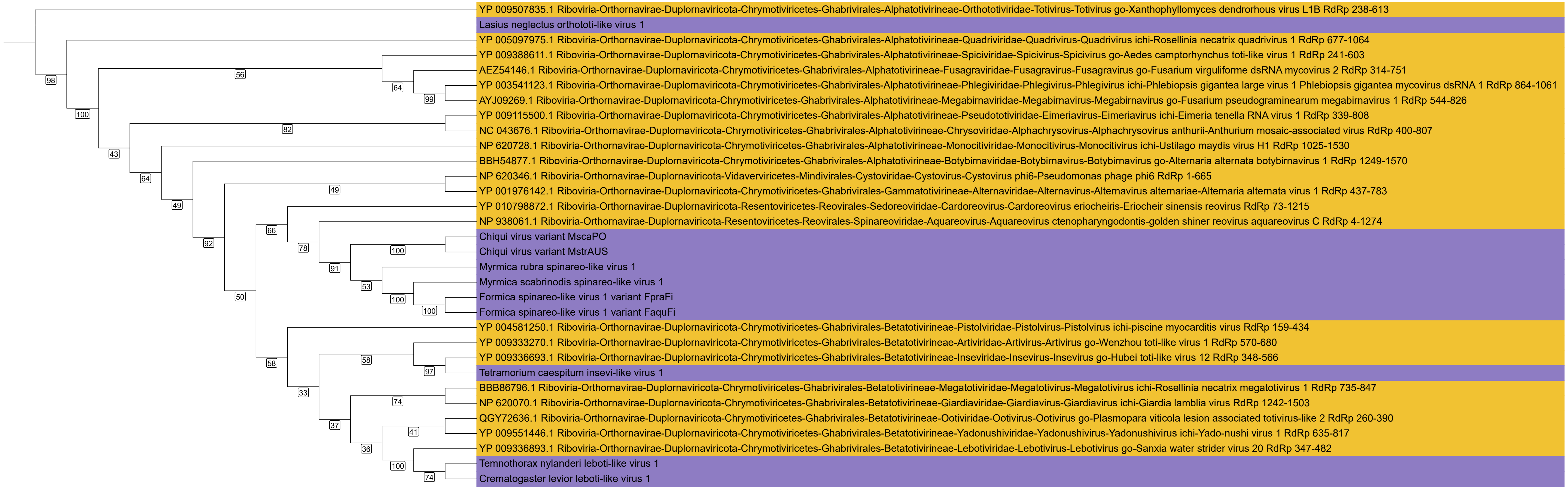

### Supplementary Figure 6. Negarnaviricota

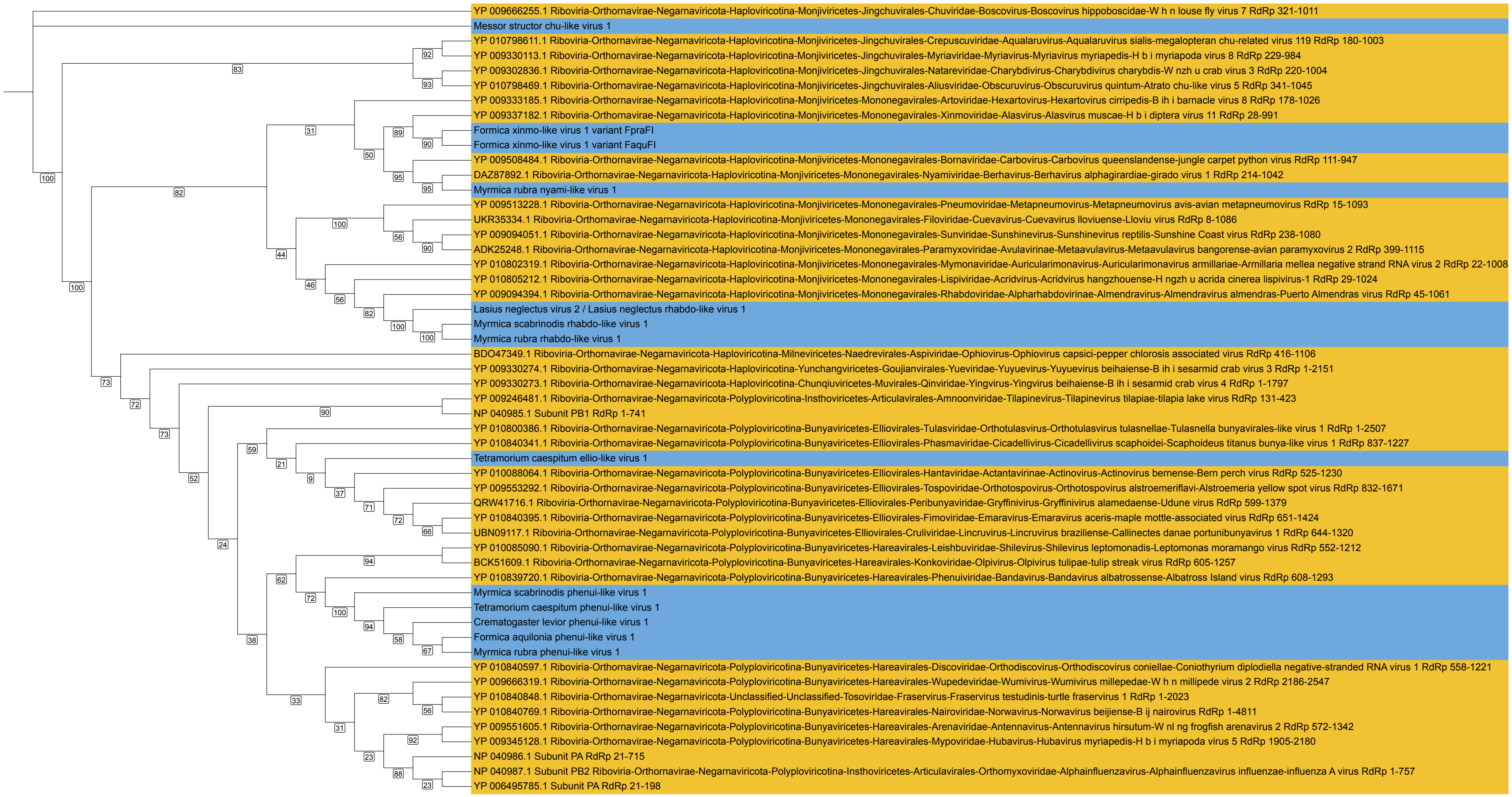
